# Cellular Anatomy of the Mouse Primary Motor Cortex

**DOI:** 10.1101/2020.10.02.323154

**Authors:** Rodrigo Muñoz-Castañeda, Brian Zingg, Katherine S. Matho, Quanxin Wang, Xiaoyin Chen, Nicholas N. Foster, Arun Narasimhan, Anan Li, Karla E. Hirokawa, Bingxing Huo, Samik Bannerjee, Laura Korobkova, Chris Sin Park, Young-Gyun Park, Michael S. Bienkowski, Uree Chon, Diek W. Wheeler, Xiangning Li, Yun Wang, Kathleen Kelly, Xu An, Sarojini M. Attili, Ian Bowman, Anastasiia Bludova, Ali Cetin, Liya Ding, Rhonda Drewes, Florence D’Orazi, Corey Elowsky, Stephan Fischer, William Galbavy, Lei Gao, Jesse Gillis, Peter A. Groblewski, Lin Gou, Joel D. Hahn, Joshua T. Hatfield, Houri Hintiryan, Jason Huang, Hideki Kondo, Xiuli Kuang, Philip Lesnar, Xu Li, Yaoyao Li, Mengkuan Lin, Lijuan Liu, Darrick Lo, Judith Mizrachi, Stephanie Mok, Maitham Naeemi, Philip R. Nicovich, Ramesh Palaniswamy, Jason Palmer, Xiaoli Qi, Elise Shen, Yu-Chi Sun, Huizhong Tao, Wayne Wakemen, Yimin Wang, Peng Xie, Shenqin Yao, Jin Yuan, Muye Zhu, Lydia Ng, Li I. Zhang, Byung Kook Lim, Michael Hawrylycz, Hui Gong, James C. Gee, Yongsoo Kim, Hanchuan Peng, Kwanghun Chuang, X William Yang, Qingming Luo, Partha P. Mitra, Anthony M. Zador, Hongkui Zeng, Giorgio A. Ascoli, Z Josh Huang, Pavel Osten, Julie A. Harris, Hong-Wei Dong

## Abstract

An essential step toward understanding brain function is to establish a cellular-resolution structural framework upon which multi-scale and multi-modal information spanning molecules, cells, circuits and systems can be integrated and interpreted. Here, through a collaborative effort from the Brain Initiative Cell Census Network (BICCN), we derive a comprehensive cell type-based description of one brain structure - the primary motor cortex upper limb area (MOp-ul) of the mouse. Applying state-of-the-art labeling, imaging, computational, and neuroinformatics tools, we delineated the MOp-ul within the Mouse Brain 3D Common Coordinate Framework (CCF). We defined over two dozen MOp-ul projection neuron (PN) types by their anterograde targets; the spatial distribution of their somata defines 11 cortical sublayers, a significant refinement of the classic notion of cortical laminar organization. We further combine multiple complementary tracing methods (classic tract tracing, cell type-based anterograde, retrograde, and transsynaptic viral tracing, high-throughput BARseq, and complete single cell reconstruction) to systematically chart cell type-based MOp input-output streams. As PNs link distant brain regions at synapses as well as host cellular gene expression, our construction of a PN type resolution MOp-ul wiring diagram will facilitate an integrated analysis of motor control circuitry across the molecular, cellular, and systems levels. This work further provides a roadmap towards a cellular resolution description of mammalian brain architecture.

## Introduction

Mechanistic understanding of brain function requires a structural framework of brain organization. The most essential feature of brain structure is an information processing network, a large set of nodes connected through sophisticated wires. Superimposed upon this anatomical infrastructure, genetic-encoded molecular machines mediate myriad cellular physiological processes, which shape neural circuit dynamics that underlie mental activities and behavior. Historically, and largely due to advances as well as constraints of available techniques, brain networks have been explored at several descending, nested levels of granularity, from gray matter regions (macroscale), to neuron types forming each gray matter region (mesoscale), individual neurons forming each cell type population (microscale), and pre- and post-synaptic elements connecting two neurons (nanoscale)^1^. Seminal studies in past decades using functional MRI and classic anatomical tracing have achieved macroscale regional connectomes in human^2^ and other mammalian brains^3–5^, providing a panoramic overview of brain organization and a global compass for further exploration^6^. However, macroscale descriptions are coarse representations and lump sum averages of functional neural circuit organization. Nerve cells are the fundamental building blocks and computation units of brain circuits that link gray matter regions via synapses, as well as the expression units of genomic information. Therefore, an essential step toward a comprehensive understanding of brain function is to establish a cellular-resolution structural framework upon which multi-scale and multi-modal information spanning molecules, cells, circuits, and systems can be registered, integrated, interpreted, and mined.

The immense number, stunning diversity, and staggering wiring complexity of nerve cells in the mammalian brain present a formidable challenge to decipher their network organization. Since the first visualization of nerve cells and formulation of the Neuron Doctrine over a century ago, a synthetic wiring diagram of the mammalian nervous system at the cell type level, or mesoscale, still remains to be accomplished. The magnitude of this problem is indicated by estimates that the mammalian brain, with its 500–1,000 gray matter regions, has on the order of at least several thousand neuron types, each receiving inputs from and projecting outputs to multiple other types^6,7^. Despite the ever-growing range of staining and imaging techniques and steady accumulation of piecemeal datasets from individual investigators in past decades, our understanding of mammalian brain circuit organization has remained grossly incomplete and inadequate.

In the past decade, major technical advances along multiple fronts have crossed key thresholds to enable cellular resolution, large scale brain circuit mapping. First, high-throughput single cell RNA sequencing began to derive a comprehensive transcriptomic cell type census in multiple brain regions. Second, systematic genetic toolkits allowed reliable experimental access to an increasingly large set of molecular-defined neuronal subpopulations and cell types. Third, continued innovations in light microscopy enabled automated high-resolution large-volume imaging of neural structures at single axon resolution across entire rodent brains, generating quantitative and comprehensive datasets on spatial organization, morphology, and input-output connectivity. Fourth, advances in computational sciences, including machine learning, allowed analysis, sharing and management of whole-brain terabyte size datasets. Fifth, novel data science and neuroinformatics tools enabled constructions of 3D digital brain atlases and the common coordinate framework (CCF) of the mouse brain, which provides a unifying registration system for large scale cross modal integration, atlasing, and quantitative analysis.

Recognizing the emerging opportunity, the establishment of the BRAIN Initiative Cell Census Network (BICCN) provides a multi-laboratory collaborative infrastructure to systematically map the anatomical organization of neuron types and chart their mesoscale connectivity in the mouse brain. Here, we present progress of the BICCN anatomy group, combining advanced expertise in cell type-targeted genetic and viral labeling tools, high resolution automated whole-brain imaging, BARseq-based projection mapping, single neuron complete reconstruction, and state-of-the-art neuroinformatic methods for CCF registration. Focusing on the upper-limb area of the mouse primary motor cortex (MOp-ul), we present 1) the delineation of the MOp-ul borders and layers within the CCF, 2) quantitative laminar distributions of projection neuron (PN) types with corresponding molecular markers, 3) whole-brain input/output organization of PN types, 4) characterization of single PN axonal trajectories and branching motifs, and 5) open access release of all associated datasets. Together, these results enable us to derive a comprehensive, PN type-based wiring diagram of the mouse MOp-ul. This will facilitate an integrated analysis of motor control infrastructure across the molecular, cellular, and systems levels, and provide a roadmap towards a mesoscale mammalian brain architecture.

## Results

We begin by establishing a cross-laboratory anatomical analysis platform that comprises: 1) classic anterograde and retrograde circuit tracing methods^3,8^; 2) genetic “driver” mouse lines expressing Cre recombinase for cell type-specific labeling^9–12^, 3) Cre-dependent AAV vectors^13–15^ and rabies virus vectors^16–18^ for anterograde and retrograde transsynaptic labeling, respectively, 4) high-throughput single-cell projection mapping using BARseq, 5) automated block-face microscopy for high-resolution mouse brain imaging: serial two-photon tomography (STPT)^19,20^, oblique light-sheet tomography (OLST)^21^, and fluorescence micro-optical sectioning tomography system (fMOST)^22^, and 6) a suite of computational and neuroinformatics approaches that includes: 3D registration to the Allen Mouse Brain Common Coordinate Framework (CCF)^23–26^; computational cell detection, axon density quantification, and neuronal morphology reconstruction across whole brain datasets^4,19,27–31^; and finally cloud-based, multi-dataset visualization at full imaging resolution^32,33^ (**Fig. 1** and Methods). Importantly, all datasets and informatics tools are presented as open resources to the neuroscience community and available at the Brain Image Library (BIL) Pittsburgh Supercomputing Center. A summary of Cre-driver lines and connectivity datasets can be also accessed in the following neuroglancer link: https://viz.neurodata.io/?json_url=https://json.neurodata.io/v1?NGStateID=LwZ24nSZk1JTHw

**Figure 1.**
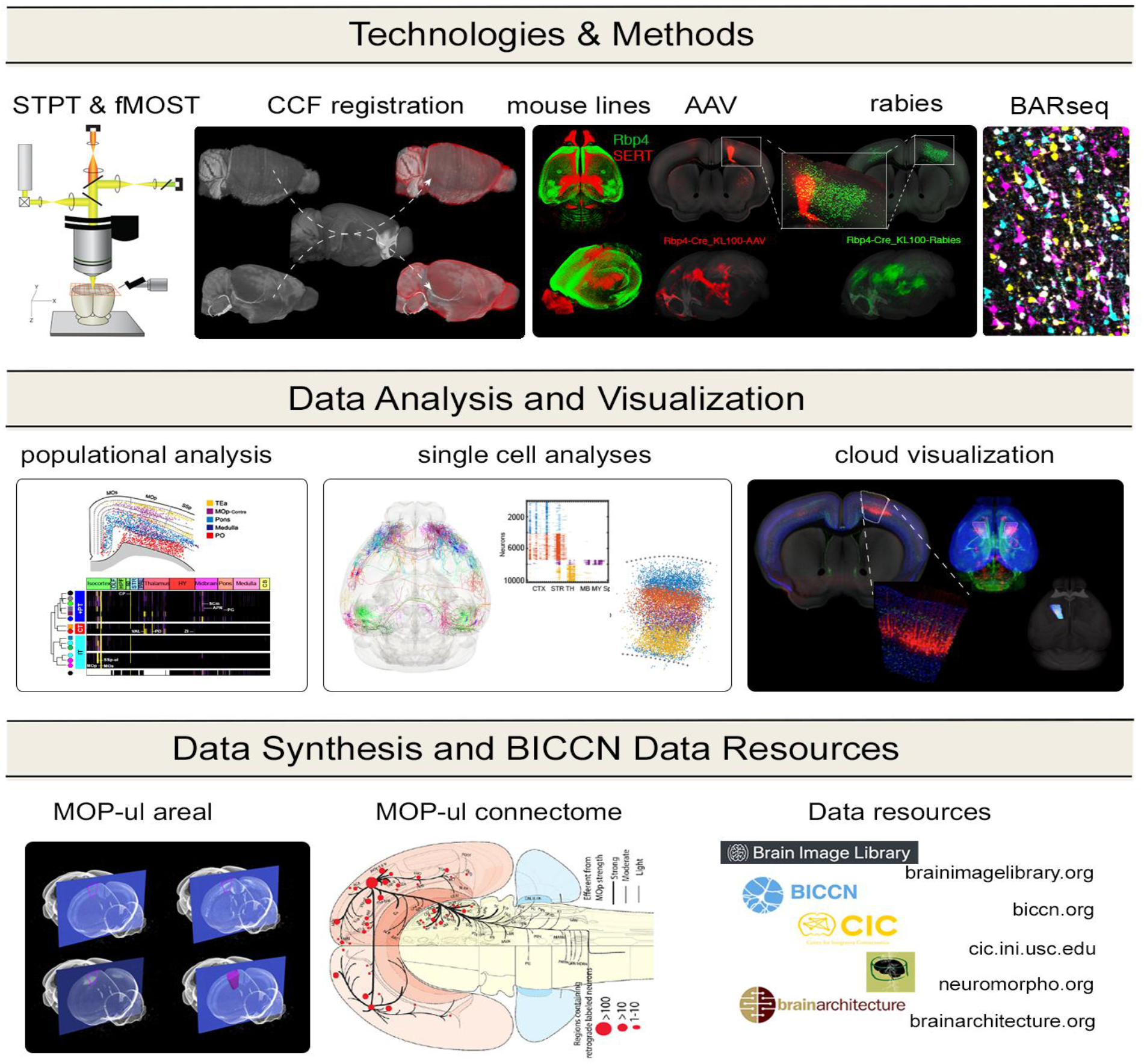
Overview of the methods, data analyses and data resources used and generated by the BICCN anatomy group. **a**, Whole brain data were generated primarily by two automated microscopy methods – STPT and fMOST. All datasets were co-registered in CCF, including cell type distribution data used for MOp-ul regional and areal delineations and transgenic lines used for projection mapping, such as the Rbp4+ transgenic line marking layer 5 neurons. Axonal projections were mapped by AAV-based anterograde tracing and BARseq, and retrograde inputs labeled with rabies-based tracing. **b**, The computational tools used in the analysis of the co-registered datasets included quantitative analyses of populational (“bulk”) labeling to map layer-specific anterograde and retrograde projections, as well as single cell morphology reconstructions and BARseq analysis to derive cell type-based single neuron projectomes. Neuroglancer was used for cloud visualization and collaborative analysis of the CCF registered data at full imaging resolution. **c**, The outcome of these efforts comprise a consensus-based delineation of anatomical borders of the MOp-ul together with a detailed description of a cortical layer- and projection neuron type-based wiring diagram.

### Delineation and cellular characterization of the MOp-ul region

MOp-ul has been traditionally localized in the rat and mouse based on cytoarchitecture, intracortical microstimulation, and connectivity tracing^34–37^, and yet no consensus has been reached regarding its spatial extent^38^, including among broadly used Paxinos and Franklin^39^, ARA^40^, and Allen CCFv3^23^ mouse brain atlases. Here we define consensus-based anatomical coordinates using a collaborative workflow, whereby multimodal image series were coregistered and cloud-visualized^32,33^ at full resolution for joint review, delineation and reconciliation of the latero-medial and rostro-caudal MOp-ul borders (**Fig. 2a**; **Supplementary Video 1**).

**Figure 2.**
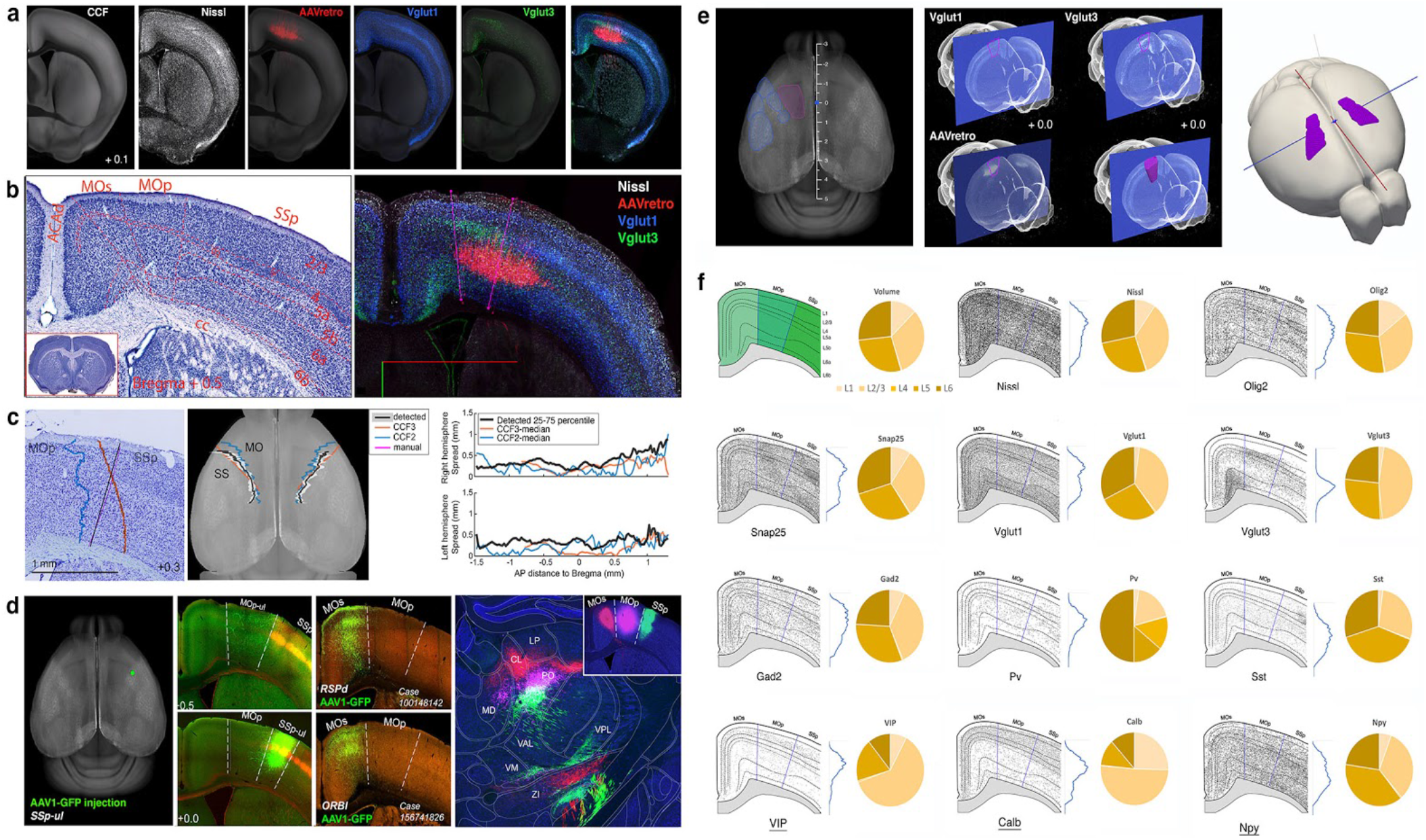
Data-driven anatomical delineation of the MOp upper limb (MOp-ul) in 2D and 3D. **a**, A total of 8 sets of anatomical imaging data in different modalities (three sets of Nissl-staining, 2 sets of viral labeling, 2 sets of cre reporter expression, and 1 set of Thy1 EGFP labeling) were co-registered in the CCF space and viewed in the Neuroglancer platform (**Supplementary Video 1**) to facilitate the delineations of the MOp-ul borders. **b**, Delineations of the MOp-ul based on Nissl-stained cytoarchitecture (left, also see **Extended data Fig. 1**), which is further enhanced and validated by areal and laminar distributions of spinal cord-projecting neurons revealed with AAV retro viral labeling after tracer injections into the cervical spinal cord, cre expression of Vglut1, Vglut3, and other cre-driver lines (**Extended data Fig. 2**). **c**, The border between MOp and SSp was algorithmically detected in 10 brains based on Nissl cytoarchitectural features to assess individual variation and relationship with CCFv2 and CCFv3 atlases. Left: Algorithmically determined boundary (black), and expert manual annotation of the MOp-SSp border (magenta) are shown together with boundaries of reference atlases diffeomorphically registered to an individual brain (CCFv2: blue, CCFv3: red). Middle: Results of the algorithmic detection mapped to the CCF space. The black lines show the median of the detected MOp-SSp boundaries with 25-75 percentile limits shown in gray. Right: The 25-75 percentile spread as a measure of dispersion (black lines) plotted together with the distances between the reference atlases and the median line (see **Extended data Figure 3** and **Supplementary Information**). **d**, Accuracy of the MOp delineation was further validated using three sets of connectivity data: (1) anterograde axonal projections from the SSp-ul to MOp-ul: transgenic mice (Scnn1a-Tg3-Cre driver line crossed with Ai14 tdTomato reporter line) received an injection of cre-dependent GFP-expressing AAV targeted precisely to the SSp upper limb area (left). Targeting restricted to the SSp was confirmed by the presence of tdTomato fluorescence in SSp layer 4. Analysis using Neuroglancer confirmed the existence of a strong monosynaptic projection from the SSp-ul to MOp-ul, therefore, confirming the border of these two adjacent cortical areas (2nd column of images); (2) the MOp medial border with the MOs was identified by the absence of a monosynaptic MOp connection with the dorsal retrosplenial area (RSPd) and ventrolateral orbital area (ORBvl) and but the presence of strong bidirectional connection between the MOs with both the ORBvl and RSPd^3^ (the 3rd column of images); (3) A triple anterograde injection strategy were used to validate global distinctive projection patterns of the MOp-ul and its adjacent SSp-ul and MOs (right). Three anterograde tracers, AAV-RFP, PHAL and AAV-GFP, were injected respectively into the newly defined MOp-ul and the adjacent MOs and SSp (Right upper corner) and their mostly non-overlapping, but topographically arranged terminal fields in the mediodorsal thalamic (MD), centrolateral (CL), paracentral (PCN) and posterior thalamic nuclei (PO). Acronyms defined in **Extended Data Table 1**. For a complete description of PHAL-labeled MOp-ul pathways please see **Supplementary Information**. **e**. The newly delineated MOp-ul was rendered into a 3D volumetric object within the CCF space. **f**, Schematic representation of MOp-ul layer delineation and laminar distribution patterns of cell populations stained by Nissl, and a subset of cre-driver lines such as pan-neuronal marker Snap25, oligodendrocytes (Olig2), main excitatory (Vglut1 and Vglut3) and inhibitory neuron markers (GAD2, Pv, Sst, VIP, Calb and Npy). Quantitative analysis of cortical-depth and layer-based distribution of the different cell types show distinct patterns of laminar distribution. Pies represent the percentage of cells per layer. Interestingly, Vglut3+ cells have prominent density on MOp-ul in contrast to MOs and SSp. See **Extended Data Fig. 8** for the quantitative analysis of all different cre-driver lines distribution.

To start, the lateral border between MOp-ul and primary somatosensory area (SSp) was derived based on the clear layer 5 transition of large MOp neuron somas versus smaller somas in the SSp cell-sparse 5a and cell-dense 5b sublayers seen in Nissl (ARA^40^; Brainmaps.org) and NeuroTrace™ stains (**Fig. 2b**; **Extended Data Figs. 1 & 2**). Next, in contrast to the classic view of the MOp as an agranular cortex, we identified a clear “granular” layer 4 of densely packed small somas continuing from SSp throughout MOp and MOs, albeit as a much narrower strip (**Fig. 2b**; **Extended Data Fig. 1**). These cytoarchitectural features also provided a robust signal for algorithmically confirming the MOp-SSp border in 10 Nissl-stained brains, revealing the extent of individual variations between animals (**Fig 2c; Extended Data Fig. 3; Supplementary Information**).

Neuronal cell type distribution and long-range projections can also assist in areal delineations^3,19,38^. The density of VGluT1+ (Slc17a7+) neurons corroborated the feature transition of layer 4 and layer 5 at the MOp/SSp border (**Fig. 2a** and **Supplementary Video 2**), while the surprisingly more restricted distribution of VGluT3+ neurons prominently highlighted the more difficult to define medial border with MOs (**Fig. 2a,b; Extended data Fig. 2**). This was further confirmed by AAV-based projection tracing from the SSp upper limb area to label the MOp-ul (**Fig. 2d** left panels), and the ventrolateral orbital area (ORBvl) and the dorsal retrosplenial area (RSPd) to label the MOs^3^ (**Fig. 2d**, middle panel). Finally, the rostro-caudal scope of the MOp-ul was defined using retrograde AAV tracing^15^ from the cervical (to delineate upper limb) or lumbar (to delineate lower limb) spinal cord (**Fig. 2a-b**; **Extended Data Fig. 4**; **Supplementary Video 1**), revealing two adjacent clusters of the upper limb domain neurons projecting to the cervical spinal cord: a medial cluster in MOp layer 5 and a lateral cluster underneath SSp layer 4 (**Fig. 2b**, right panel; **Extended data Figure 4**). The MOp-ul borders were finally validated using a triple anterograde labeling strategy – injecting AAV-RFP, PHAL, and AAV-GFP into MOs, MOp-ul, and SSp, respectively – which revealed topographically organized projection patterns that agreed with the expected corresponding origins, including mostly non-overlapping terminal fields in different thalamic nuclei and zona incerta (**Fig. 2d**; **Extended Data Fig. 5e**), MOp-ul projections to the intermediate and ventral horn of cervical spinal cord (**Fig. 2d**; **Extended Data Fig. 5l**) contrasting to SSp projections to the dorsal horn.

To facilitate further analyses, the MOp-ul borders in CCF and Neuroglancer were used to render a 3D volumetric MOp-ul template (**Fig. 2e**; **Supplementary Video 2**) aligned with the recently developed LSFM-based 3D histological data (**Extended data Fig. 6**). To compare our MOp-ul boundaries to existing atlases, we imported Allen reference atlas^40^ and Franklin-Paxinos atlas^39^ onto the Allen CCFv3^23^ (**Extended data Fig. 7**). Finally, using the STPT imaging pipeline, we quantified the cellular and cell type distribution in the MOp-ul, including glutamatergic (VGluT1+) versus GABAergic (GAD2+) neurons, major GABAergic inhibitory neuronal subpopulations, and a number of other Cre line expressions in the MOp-ul and across different layers (**Fig. 2f; Extended data Fig. 8**).

### MOp laminar organization delineated by the distribution of projection neuron types

Cortical areas comprise three broad excitatory projection neuron (PN) classes based on their projections across major brain divisions: IT (intratelencephalic) primarily targeting cortical and striatal regions, PT (pyramidal tract) or ET (extratelencephalic) primarily projecting to the lower brainstem and spinal cord, and CT (corticothalamic) projecting specifically to the thalamus^41–43^. To further examine the PN laminar distribution in MOp-ul, we performed a series of retrograde tracing experiments using classic tracers (fluorogold, FG and cholera toxin B subunit, CTB) and rabies viral tracers injected to: cerebral cortex and contralateral caudoputamen (CP) for IT neurons; thalamus for CT neurons; midbrain, pons, medulla, and spinal cord for ET neurons. In addition to confirming the general class distinctions, these experiments revealed a novel and more refined laminar distribution pattern of PNs, suggesting more than 25 PN subtypes (**Fig. 3a-c**) across eleven distinct MOp-ul layers and sublayers (1, 2, 3, 4, 5a, 5b-superficial, 5b-middle, 5b-deep, 6a-superfical, 6a-deep, and 6b).

**Figure 3.**
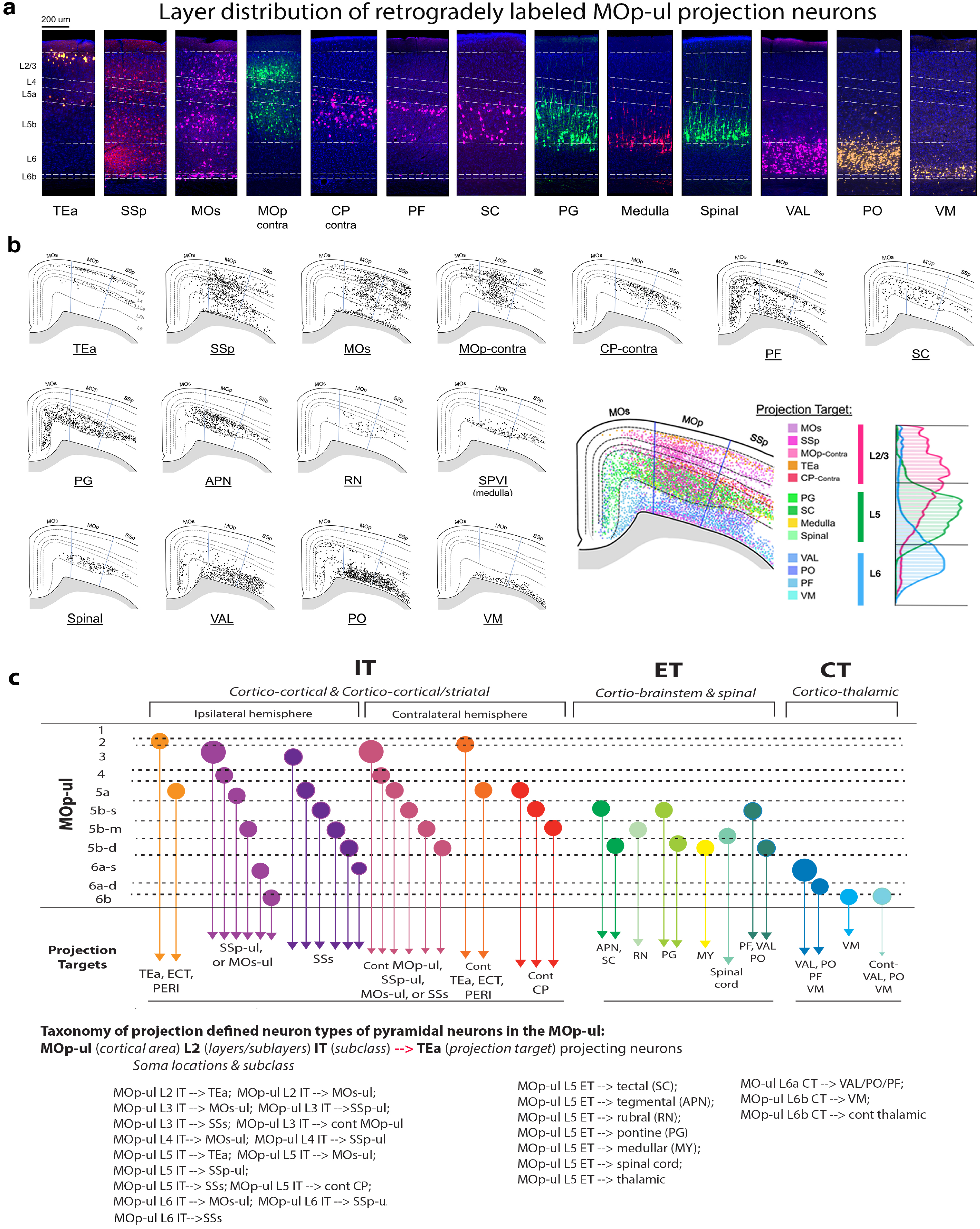
Projection target defined MOp-ul neuron types. **a**, Laminar distributions of retrogradely labeled projection neurons in the MOp-ul after tracer injections into different MOp-ul projection targets. TEa projecting neurons are distributed in layer 2 and 5a; SSp and MOs projecting neurons are distributed throughout layers 2-6b; contralateral MOp projecting neurons (commissural neurons) are in layer 3, 5a and 5b; contralateral CP projecting neurons are specifically distributed in layer 5a and 5b. All of these neurons belong to the intratelencephalic (IT) neuron type. Several PT (pyramidal-tract neurons) or ET (extratelencephalic) type neurons with projections to the PF, SC, pons, medulla, and spinal cord display sublaminar specificities in layer 5b, namely the superficial (L5b-s), middle (L5b-m) and deep (L5b-d). Finally, all corticothalamic projecting neurons (CT) are distributed in layer 6a (VAL- and PO-projecting neurons) and 6b (VM-projecting neurons). **b**, Detailed distribution patterns of those retrogradely labeled projection neurons in the MOp. Semi-quantitative analysis shows distinct laminar specificities of different neuron types with distinct projection targets. Please also see **Extended Data Figs. 10 & 12** for additional retrograde labeling in bilateral MOp. **c**, A schematic representation of our anatomical classification of cortical projection neuron types based on their soma locations and projection targets (based on retrograde labeling). These neuron types are named according to their laminar specificities (L2-L6), projection neuron subclasses (IT, PT and CT), and their projection targets. Collateral projections of these neuron types were further characterized using other methods (see next sections). Acronyms defined in **Extended Data Table 1**.

As expected, IT neurons are distributed broadly in layers (L) 2-6b (**Fig. 3a-c; also see Extended data Figs. 9 & 10**). Individual layers contain intermingled IT neurons innervating different targets, while neurons targeting the same structures can be distributed in different layers and display distinguishable connectivity properties. Several IT types were identified: 1) two TEa-projecting types: a L2 type that generates an asymmetric projection pattern with denser innervation to the contralateral TEa, and a L5a type that projects relatively symmetrically to bilateral TEa (also see **Extended data Fig. 9**). 2) IT neurons that target other somatic sensorimotor areas (e.g. MOs-ul, SSp-ul and SSs) in the ipsilateral hemisphere are intermingled in layers 2, 3, 4, 5a, 5b-middle sublayer of the MOp-ul. These layers mainly contain PNs that project to the contralateral MOp-ul, while those projecting to the contralateral MOs, SSp and SSs are relatively sparse (also see **Extended data Fig. 10**). 3) L4 contains many MOs- and SSp-projecting neurons, but much less SSs-projecting neurons. 4) L6b also contains a dense cluster of IT neurons that project to the ipsilateral but not contralateral MOs and SSp (also see **Extended data Fig. 10**). Very few L6b neurons project to SSs. Finally, cortico-striatal projecting IT neurons are distributed preferentially in layers 5a and 5b superficial and middle sublayers (also see **Extended data Figure 10**).

ET (also known as PT) neurons are distributed primarily in L5b ^42,43^. A preferential superficial and deep sublayer distribution of retrograde labeling in L5b was observed after injections into certain regions of the thalamus (parafascicular nucleus), midbrain (anterior pretectal nucleus and superior colliculus), and hindbrain (pontine nuclei); whereas a preferential middle to deep sublayer 5b distribution of retrograde labelling was observed after injections into other regions of the midbrain (red nucleus), the medulla (e.g. spinal nucleus of trigeminal nerve interpolar part), and cervical spinal cord, with the deepest L5b labeling resulting from medulla injections (**Fig. 3a-c**).

Among L6 CT neurons, we observed substantial differences between L6a and L6b. L6a neurons primarily project to the posterior thalamic complex (PO), ventral anterior-lateral thalamic complex (VAL), and parafascicular nucleus (PF), as well as the reticular thalamus (RT). Another type of CT neurons generating primary projections to the ventromedial thalamic nuclues (VM) are distributed in L6b and adjacent deep L6a. Finally, we also identified a specific type of L6b CT neurons that specifically project to the contralateral thalamic nuclei, such as the PO, VAL and VM (**Figure 3a-c; Extended Data Fig. 10**).

Next, we systematically examined spatial distribution of 40 genes selected from the Allen Brain Atlas gene expression database to recapitulate laminar specificities of different PN types in the MOp-ul (**Extended data Fig. 11**). Genes distributed specifically in single layers, thus, providing putative candidate markers for specific PN types, include (1) Trpc6 for L2 IT; (2) Nr5a1-cre and Scnn1 for L4 IT; (3) Tnnc1 and Trib2 for L5a IT; and (4) Tle4, Ntsr1-cre and Sulf1 for L6 CT. Some genes recapitulate the sublaminar specificities of L5b, for example, Sim1-cre, Efr3a-cre, Hsd11b1 and Chrna2-cre in L5b superficial sublayer; and Postn, Npr3-cre, Gng7, and Layn in L5 middle and deep sublayers. Other genes are expressed in multiple layers and presumably intermingled PN types, such as Cux2-cre (L2/3/4 IT), Plxnd1-cre (L2/L5a IT), Tlx3-cre (L5a/5b IT), and Foxp2 (L5/6 IT/PT/CT). Several L6b-specific genes include Ctgf, Cplx3, Nxph4 and Trh. But, it remains unclear whether they are L6 IT or CT neurons. Retrograde tracing in combination with spatial transcriptomics (e.g. MERFISH^44^) is necessary to validate these molecular markers for different PN types.

Overall these results reveal a wide variety of MOp-ul projection neurons that can be initially classified according to their sublaminal origin and projection targets (**Fig. 3c**). We have adopted a straightforward scheme to represent these PN types, For example, MOp-ul L2→TEa for MOp-ul layer 2 neurons projecting to TEa (see **Fig. 3c** for a complete list of PN types based on these results). Whole brain axonal projections of these neuron types were further characterized using several other methods described below.

### MOp-ul output patterns

We systematically examined neural outputs of the MOp-ul at two different resolutions: 1) using the classic PHAL tracer ^3,8^, we first characterized the overall MOp-ul output patterns. Axonal pathways arising from the MOp-ul region project to over 110 targets in the brain and spinal cord, with 60 receiving moderate to dense innervation (**Extended Data Figs. 5, 12; Extended Data Table 2; Supplementary Information**); 2) using Cre-dependent viral tracers with injections into different Cre lines selective for these laminar- and projection-cell subclasses^4,45^, we examined projections from L2/3 IT, L4 IT, L5 IT, L5 ET, and L6 CT cells (**Fig. 4a-c**). As these two methods label axons including presynaptic terminals, synaptic innervation (versus fibers of passage) within the target areas was also confirmed using two different viral tracing methods (**Extended Data Fig. 13**; see also^45^).

**Figure 4.**
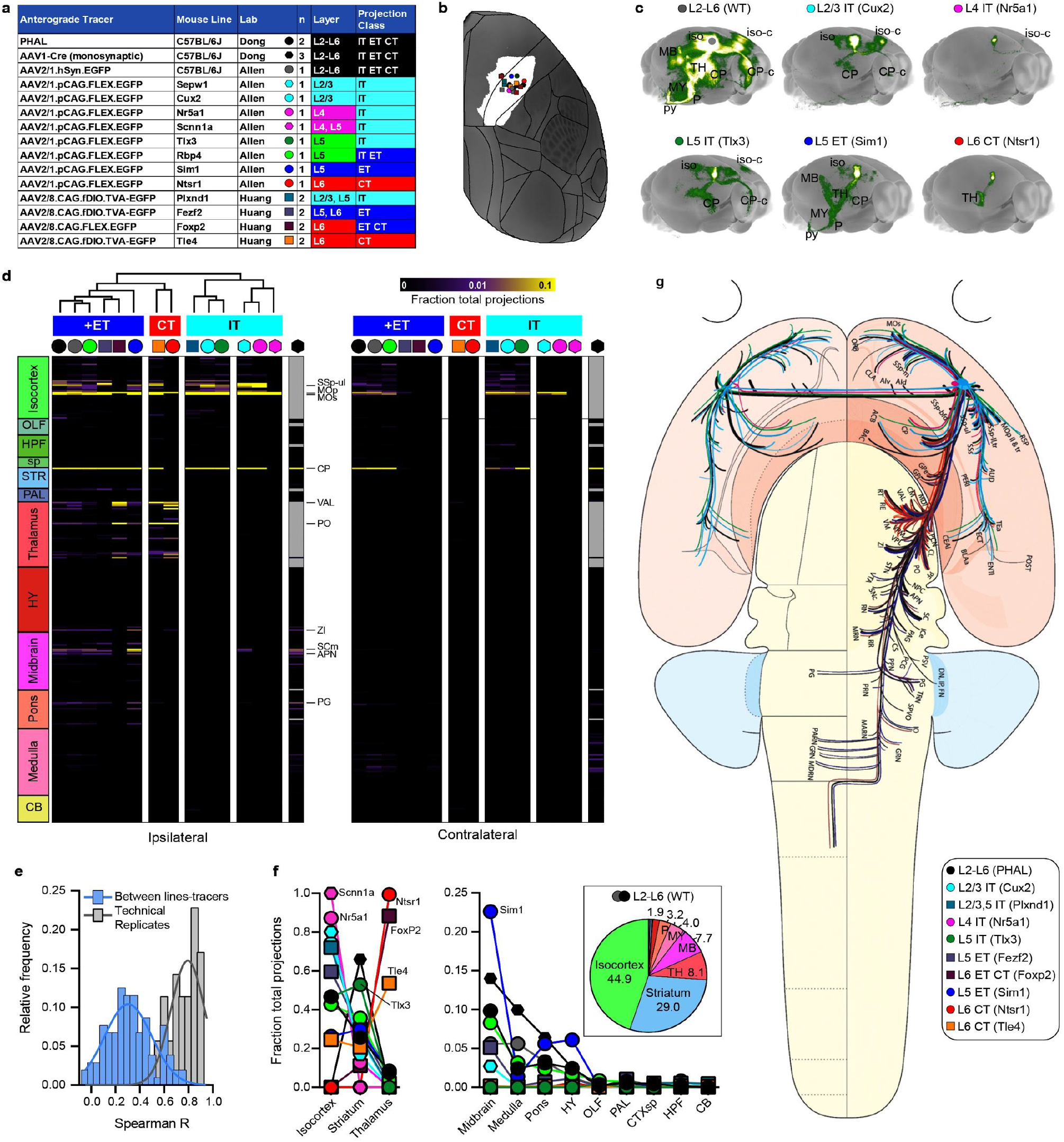
Brain-wide MOp-ul projection patterns by layer and class. **a**, Key shows the different tracers and mouse lines used across three labs to label axons originating from neurons in primary motor cortex. The layer and projection class selectivity for each Cre driver line is also summarized. Symbols and color code are used in b-g. **b**, Injection locations are plotted on a top-down view of the right cortical hemisphere from CCFv3 with the MOp-ul delineation from **Fig. 2** shown in white. The mean distance and SD between all injection centroids is 443.0 +/- 185.04 μm. **c**, Maximum intensity projections show the brain-wide distribution of labeled axons from all classes in MOp-ul (top left), and from distinct layer and class defined by Cre lines. **d**, A directed, weighted connectivity matrix (15 x 628) from MOp to 314 ipsilateral and 314 contralateral targets for each of the fifteen mouse lines or tracers listed in (a). Each row shows the fraction of the total axon signal measured from a single experiment or the average when n>1. Signal in the injection site (MOp) was subtracted from the whole brain total to show the fraction of projections outside of the injected region. Columns are ordered by major brain division. For AAV1-Cre monosynaptic tracing, known reciprocally connected regions were excluded from analyses and are colored gray. We performed hierarchical clustering with Spearman rank correlations and complete linkages, splitting the resulting dendrogram into four clusters. One contained all experiments, across all labs, in which ET neurons were labeled. The next cluster contained CT lines (Tle4 and Ntsr1). The third and fourth clusters contained all the IT lines. We did not include AAV1-Cre in the clustering due to the many excluded regions. Selected target regions are indicated with a line and acronym. The color map ranges from 0 to 0.1; it is truncated at the top. **e**, Frequency distributions of Spearman’s correlation coefficients (R) from the dataset in (d) and for Rs measured between individual experimental replicates in MOp. A curve was fit to each distribution (lines). The distribution of Spearman Rs between different line-tracer experiments is normally distributed with weaker correlations than for the replicates (mean = 0.30 v 0.79). **f**, The fraction of total projections is plotted for each line/tracer across 12 major brain divisions. The pie chart inset shows the % of total axons in the PHAL and AAV experiments in WT mice. Most projections from MOp-ul target regions within isocortex, striatum, thalamus and midbrain, with relatively fewer projections to the medulla and pons. The fraction of total projections in each major division reflect the projection class labeled by different Cre lines. **g**, Schematic summarizes all major MOp outputs by area, layer, and projection class on a whole brain flatmap for anatomical context (please see **Extended Data Fig. 15** for individual flatmaps of projection pathways revealed with different cre line/tracer experiments). Acronyms defined in **Extended Data Table 1**.

Axon signals from all anterograde tracer experiments were mapped to CCFv3 for quantitative comparison across 314 non-overlapping ipsilateral and contralateral gray matter regions^23^. We generated a directed and weighted connectivity matrix with edges defined as the proportion of axon per target region, (**Fig. 4d; Extended Data Table 2**)^4,45^. Notably, Cre line-labeled projections classes revealed highly distinct components of the overall output pathway revealed by PHAL and AAV (**Fig. 4c,d, Extended Data Figs. 5, 12, 14; also see Supplementary Information**). The projections of Sepw1–L2/3, Cux2–L2/3, Nr5a1–L4, Scnn1a–L4/5, Plxnd1– L2/3+L5, and Tlx3–L5 were restricted to the isocortex and CP, the defining IT feature. The projections of Sim1–L5 and Fezf2–L5/6 were predominantly distributed subcortically, consistent with the ET classification. The projections of Ntsr1–L6 and Tle4–L6 targeted thalamic nuclei, reflective of CT neuron types (**Fig. 4c,d**). Several Cre lines labeled multiple PN classes, including IT and ET in Rbp4–L5 (**Fig. 4c,d**), and PT and CT in Foxp2–L6 (**Fig. 4d**).

To compare all tracing experiments, we performed unsupervised hierarchical clustering based on the **Fig. 4d** connectivity weights, cutting the dendrogram into four main clusters. Cluster 1 comprised all L5 ET projection classes, including the global patterns derived from PHAL and AAV-GFP and the Rbp4–L5 mixed IT/ET labeling. Cluster 2 comprised the Ntsr1–L6 and Tle4-L6 CT projections. Clusters 3 and 4 contained all IT projections, with cluster 3 comprising Cux2–L2/3, Tlx3-L5, and Plxnd1–L2/3+L5, and Cluster 4 comprising Sepw1–L2/3, Nr5a1–L4, and Scnn1a–L4 (**Fig. 4d**). Notably, the correlations between projection patterns for all pairs of lines and tracers (columns in **Fig. 4d**) were significantly lower than for technical replicates, indicating biological rather than technical variability (p<0.0001, Mann-Whitney test, **Fig. 4e**).

The overall output from MOp-ul is predominantly to the isocortex, striatum, and thalamus (44.9, 29.0, and 8.1% of total axon density, respectively) with relatively less axons detected in the midbrain, medulla, and pons (**Fig. 4f**). Our result includes several previously unreported target areas (also see **Extended data Fig. 14** and **Supplementary Information**), including capsular central amygdalar nucleus (CEAc) in the striatum, bed nucleus of the anterior commissure (BAC), globus pallidus external segment (GPe), contralateral thalamic nuclei (PCN), and cerebellar interposed nucleus (IP). Although our clustering methods primarily confirmed the visual classification of anterograde tracing experiments into expected major projection classes based on differential targeting of major brain divisions, we also observed differences between lines from the same class/cluster. At the PN subpopulation level, most labeled axons in IT lines were in isocortex and striatum, but their relative distributions differed by Cre line (**Fig. 4f**, left). Specifically, the L4 IT lines (Scnn1a, Nr5a1) had more axon in isocortex compared to both L2/3 and L5 IT lines (1.0 and 0.87 *vs*. 0.8, 0.74, 0.72, 0.47; Sepw1–L2/3, Cux2–L2/3, Plxnd1– L2/3+L5, and Tlx3-L5, respectively). In contrast, Tlx3–L5 had a much larger fraction of projections into striatum compared to the other IT lines (0.53 *vs*. 0.17, 0.25, and 0.26; Sepw1– L2/3, Cux2–L2/3, and Plxnd1–L2/3+L5, respectively). The two Cluster 2 L6 CT lines also displayed clear specificity differences: while nearly 100% of axons from Ntsr1–L6 terminated in the thalamus and primarily in the ventral anterior lateral nucleus (VAL), only 54% of the Tle4-L6 axons terminated in the thalamus, with most of the remaining axons in isocortex and striatum. Within the thalamus, Ntsr1 neurons generate dense projections to the PF and RT, which receive rather sparse input from Tle4 neurons.

Further analysis revealed the predominant PN types constituting different MOp-ul output channels (**Extended Data Fig. 14**). For example, projections to SSp-ul originate from both L2/3 and L5 IT neurons labeled in the Cux2–L2/3, Rbp4–L5 and Tlx3-L5 populations, with little-to-no contribution from neurons labeled in the Sim1–L5 ET, and Ntsr1–L6 CT lines. Similarly, projections to CEAl are from L5 IT cells (Rbp4-L5 and Tlx3-L5, but not Cux2-L2/3, Sim1-L5, or Ntsr1-L6, **Extended Data Fig. 14**), and projections to GPe are primarily from L5 ET cells (Rbp4, Fezf2, Foxp2, Sim1). These experiments also confirmed (in the Cux2-L2/3 IT line) the unique population of L2 neurons projecting contralaterally to TEa, ECT, and PERI identified by retrograde tracing (**Extended Data Fig. 14**), and the Tle4-L6 CT projection to contralateral thalamic nuclei. Moreover, the sparse cerebellar projection to the IP nucleus, also detected in the AAV1-Cre monosynaptic tracing result (**Extended Data Table 2**), is labeled in Rbp4-L5 but not in other L5 ET lines. This suggests that current Cre drivers may not sufficiently label the population of cerebellar projecting neurons seen in both PHAL and AAV-GFP tracing. Altogether, we revealed a PN type-based output stream of the MOp-ul (**Fig. 4g**, and **Extended Data Fig. 15**).

### MOp-ul input pattern

To map brain-wide inputs to MOp, we combined several different tracing experiments (**Fig. 5a-c**): 1) classic retrograde tracing with cholera toxin subunit B (CTB) (also see **Extended data Fig. 12**); 2) Cre-dependent monosynaptic rabies viral tracing in the above described Cre lines targeting major IT, L5 ET, and CT PN types, plus three additional interneuron-selective lines (Pvalb-, Sst-, Vip-Cre); and 3) a modified TRIO (tracing the relationship between input and output) strategy combining AAV-retro Cre with monosynaptic rabies viral tracing from distinct projection-defined neuron types^46^ (**Extended Data Fig. 16**).

**Figure 5.**
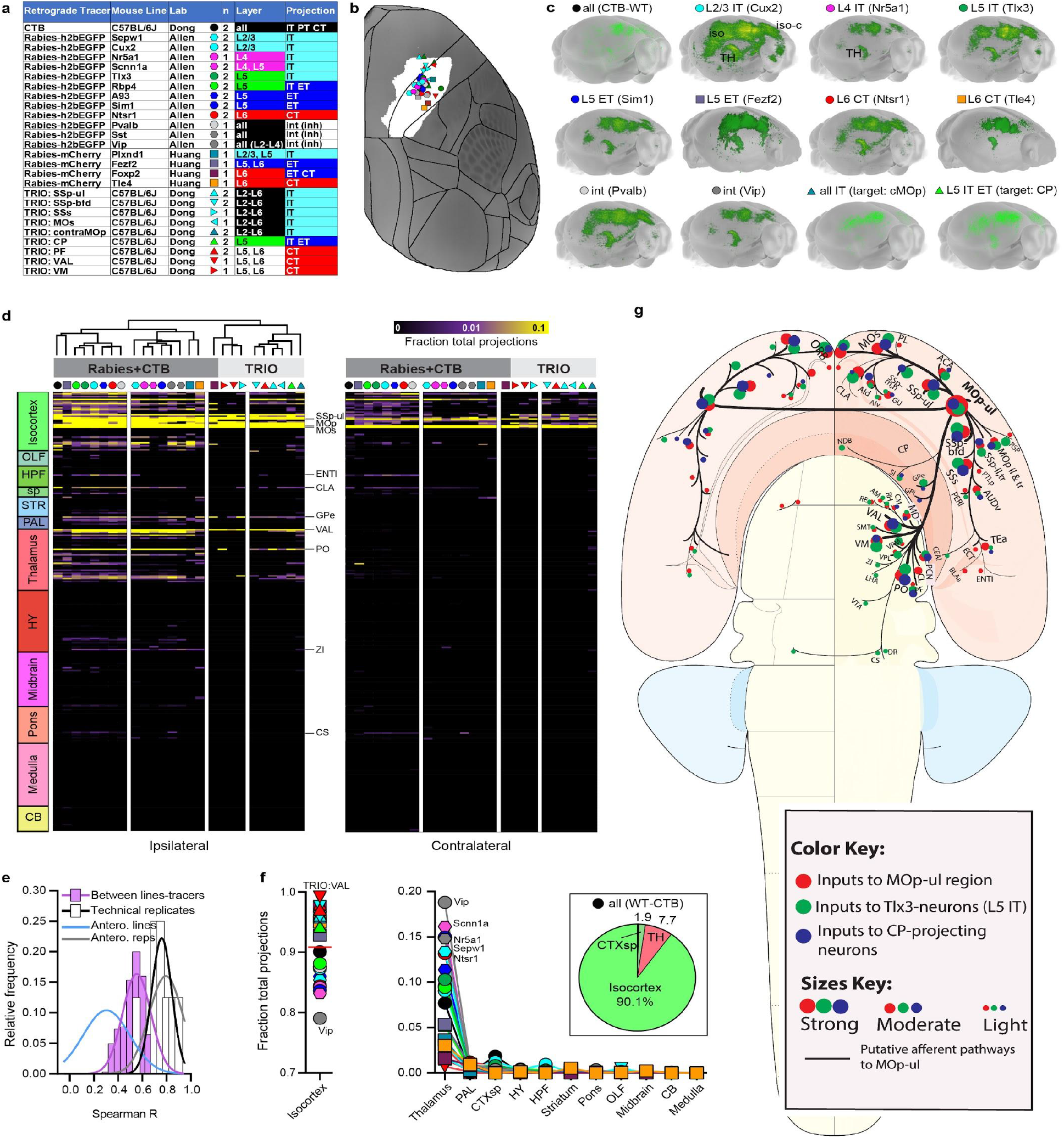
Brain-wide inputs to MOp-ul by layer and class. **a**, Key shows the different tracers and mouse lines used across three labs to label inputs to cells in the MOp-ul. The layer and projection class selectivity for each Cre driver line or tracer type is also summarized. Symbols and color code are used in **b-g**. **b**, Injection locations are plotted on a top-down view of the right cortical hemisphere with the MOp-ul delineation from **Fig. 2** shown in white. The mean distance and SD between all injection centroids is 622.4 +/- 337.01 μm. **c**, Maximum intensity projections show the brain-wide distribution of inputs to MOp, by distinct layer or cell class defined by Cre lines or TRIO. Note the strong similarities in patterns for all experiments. **d**, A directed, weighted connectivity matrix (26 x 628) to MOp from 314 ipsilateral and 314 contralateral targets for each of the mouse lines or tracers listed in **a**. Each row shows the fraction of the total input signal measured from a single experiment or the average when n>1. Signal in the injection site (MOp) was subtracted from the whole brain total to show a fraction of inputs outside of the injected region. Columns are ordered by major brain division. Fraction of total input across these 628 structures was used to assess for similarities and differences between lines, tracers, and labs. Following hierarchical clustering with Spearman rank correlation and complete linkage we split the dendrogram into two major clusters. The first split contains all rabies and CTB experiments, with the exception of rabies tracing from MOp in the FoxP2-Cre line. The second split contains all TRIO experiments. Within these two major clusters, subclusters can be differentiated based on the amount of subcortical input (for rabies and CTB) or cortical input (for TRIO). Selected input regions are indicated with a line and acronym. The color map ranges from 0 to 0.1; it is truncated at the top. **e**, Frequency distributions of Spearman’s correlation coefficients (R) from the dataset in **d** and for R measured between individual experimental replicates in MOp. A curve was fit to each distribution (lines). Anterograde experiment curve fits from **Fig. 4** are included for comparison. The mean of the distribution of Spearman Rs between retrograde tracing experiments is notably closer to the replicate mean (0.55 v 0.76) compared to the anterograde tracing experiments. **f**, The fraction of total inputs is plotted for each line/tracer across 12 major brain divisions. The pie chart inset shows the % of total inputs from the CTB experiment in WT mice to summarize the total brain-wide distribution across all layers/classes. Most input to MOp-ul is from regions within isocortex, followed by thalamus across all lines/tracers.**g**, Schematic summarizes (major) MOp inputs by area (red), layer (i.e., L5 IT Tlx3 neurons, green), and projection class (i.e., CP-projecting neurons, blue) on a whole brain flat map for anatomical context. The sizes of dots represent relative strength of connectivity. Acronyms defined in **Extended Data Table 1**.

CTB tracing revealed a broad set of input source areas to MOp-ul, including somatomotor cortical regions (MOp, SSp, SSs, MOs) and related thalamic components (VAL, PF, PO and VM) (**Extended data Figs. 12, 16**). In contrast to projection mapping results, rabies tracing from Cre line- and target-defined neuron classes revealed overall similar global inputs sources to those observed with CTB (**Fig. 5c, Extended Data Figs. 16, 17**). One notable difference is that rabies viral tracing revealed additional inputs to MOp from pallidal regions (GPe, GPi, and CEAc) and from putative monoaminergic inputs from brainstem, such as the compact part of substantia nigra (SNc), superior central nucleus raphe (CS), and dorsal raphe (DR) (**Fig. 5c, Extended Data Fig. 16**). We quantified the fraction of total labeled inputs in each of the 314 ipsilateral and contralateral gray matter regions and generated a directed, weighted, input connectivity matrix (**Fig. 5d**, see also Methods and **Extended Data Table 3**). We then performed unsupervised hierarchical clustering of the brain-wide input patterns from all 26 distinct retrograde tracing experiments (**Fig. 5d**). Unlike for the MOp-ul output datasets, the resulting dendrogram revealed highly similar input connectivity patterns across projection classes, with the two main clusters associated with tracing methods. The first larger cluster comprised CTB and Cre line-based datasets. The second cluster comprised all TRIO-based experiments and the Cre-dependent tracing from the Foxp2–L6 line. The in-degree for these two major clusters was significantly different (avg. n=91 vs 30 sources, p<0.0001, two-tailed t-test), suggesting on average narrower labeling from the TRIO experiments. Although the correlations between input patterns for all pairs of lines and tracers (columns in **Fig. 5d**) were significantly lower than for technical replicates (p=0.02, Kruskal-Wallis test with Dunn’s multiple comparison post-hoc), the mean correlation was significantly higher for retrograde vs. anterograde experiments across similar sets of Cre lines (avg R=0.57 vs 0.33, **Fig. 5e** purple and blue lines). Overall, input to MOp-ul is predominantly from isocortex and thalamus, with fewer input cells in subcortical areas such as the cortical subplate (**Fig. 5f**).

Further comparison of rabies labeling from Cre-defined cell classes and TRIO to CTB tracing confirmed highly similar distributions of monosynaptic inputs (**Fig. 5d**; **Extended Data Figs**. **16, 17**), with robust labeling in SSp-ul L2/3 and L5 (**Extended Data Fig. 16**, second row), ipsilateral thalamic VAL, and claustrum. Altogether, these data suggest that input sources to Cre- and target-defined MOp-ul neuron populations are similar, consistent with other recent findings that global input patterns are independent of starter cell type^47–49^. Nevertheless, axonal inputs to the MOp-ul arising from different cortical and thalamic regions display clear laminar preferences (**Extended data Fig. 18**) consistent with previous literature^50^. These results also do not exclude the possibility that distinct presynaptic neuron types within the same source area may project to different PN or interneuron types within the MOp. Schematically analyzing these input sources together with the MOp output targets (**Fig. 5g; Extended Data Fig. 12**) demonstrates that all input sources to MOp were also its projection targets, indicating prevalent reciprocal areal connections with mostly compatible inputs/outputs strengths (**Extended Data Fig. 12**).

### High-throughput projection mapping with single-cell resolution

While driver line-dependent and target-defined tracing resolves PNs to distinct subpopulations, they do not achieve single cell resolution and require large numbers of injections in many individual animals. The BARseq method achieves high-throughput projection mapping at cellular resolution using *in situ* sequencing of RNA barcodes^51^. Using BARseq^51^, we mapped the projection patterns of 10,299 MOp neurons to 39 target brain areas (**Fig. 6a**). MOp projection patterns obtained by BARseq were comparable to those obtained by single-cell tracing (See **Supplementary Information**; **Extended Data Fig. 19a-f**), and the large sample size from BARseq can reveal additional statistical structures in projections (See **Supplementary Information**; **Extended Data Fig. 19g-i**).

**Figure 6.**
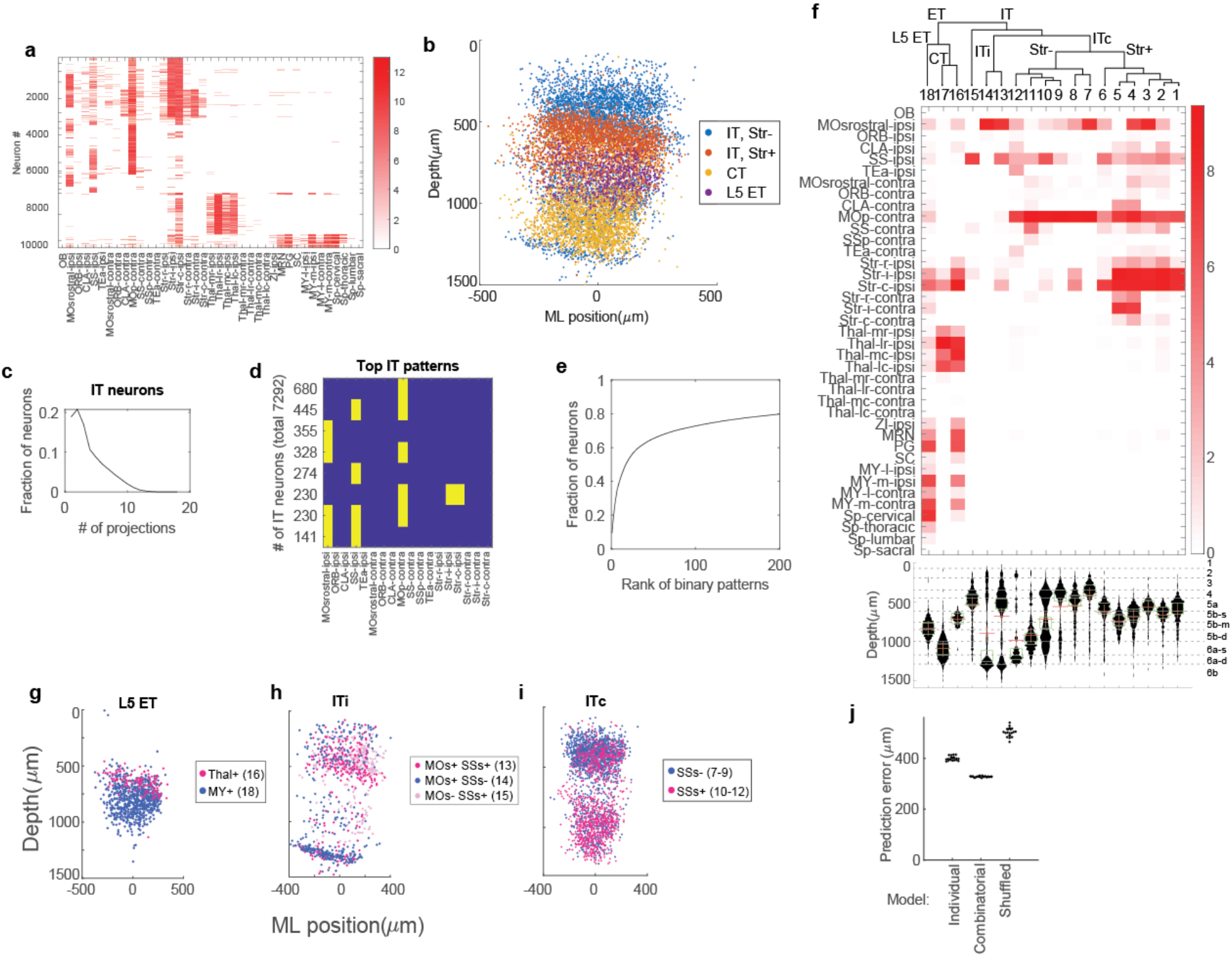
Projection mapping with single cell resolution using BARseq. **a**, Log transformed projection patterns of 10,299 neurons mapped in the motor cortex. Rows indicate single neurons and columns indicate projection areas. **See Supplementary Table 1** for a detailed list of dissection areas. **b**, Scatter plot of soma locations of the mapped neurons in the cortex. X-axis indicates relative medial-lateral positions, and y-axis indicates laminar depth. Neurons are colored by major classes as indicated. **c**, Distribution of projection numbers of IT neurons. **d**, Top binary projection patterns of IT neurons. The number of neurons with each pattern is indicated on the left. **e**, Cumulative fractions of IT neurons (y-axis) with the indicated number of binary projection patterns (x-axis). The projection patterns are sorted by their abundances, so the most common patterns are on the left. **f**, Mean projection strengths of the indicated subgroups. Rows indicate projection areas and columns indicate subgroups. Dendrogram constructed from the distance of mean projection patterns is shown above, with major classes and splits indicated. Histograms of the laminar distribution of subgroups are shown below. In the histograms, sublayer identities as defined in **Fig. 3c** are indicated on the right, and sublayer boundaries are indicated by dashed lines. **g-i**, Scatter plot of soma locations of the indicated subgroups in the cortex. X-axes indicate relative medial-lateral positions, and y-axes indicate laminar depth. Group numbers are shown in parentheses. Major classes to which the neurons shown belonged to are indicated above each panel. **j**, Mean prediction error of laminar depths using models trained on individual projections, combinatorial projections, and shuffled projections. p < 4 ×10^−24^ for all pairwise comparisons using two-sample t-tests.

Hierarchical clustering revealed the three major classes of projection neurons (CT, L5 ET, IT), as well as two sub-classes of IT neurons, those with or without projections to the striatum (IT Str- or cortico-cortical and IT Str+ or cortico-striatal). Consistent with previous reports and the tract tracing results aforementioned, these four classes of neurons occupy distinct laminar positions (**Fig. 6b**). Most (81%) IT neurons projected to two or more areas (**Fig. 6c**) with some neurons (1.9% of neurons) projecting to 11 or more areas. The most common binary projection patterns were dominated by combinations of projections to ipsilateral MOs, SSs, and contralateral MOp, consisting of seven out of the top eight binary projection patterns (**Fig. 6d**) and 34% of all IT neurons (**Fig. 6e**). Because our analysis would miss projections to areas that BARseq did not sample, our estimates of projection numbers per neuron are likely a lower bound.

Within each class, further divisions by projection patterns (see **Methods**) resulted in 18 subgroups with distinct laminar distributions (**Fig. 6f**). These subgroups revealed several combinatorial projection patterns beyond the major class-level divisions that were enriched in specific sublayers as defined in **Fig. 3c**. For example, we found two types of ET neurons (groups 16 and 18) with distinct laminar positions, one predominantly projecting to the medulla ventral to one projecting to the thalamus (**Fig. 6g; Extended Data Fig. 19j**). This result is consistent with our retrograde tract tracing results (**Fig. 3a, b**), which shows PF-projecting neurons in L5b-superfical sublayer while medulla-projecting neurons in L5b-deep sublayer. Such divisions in ET neurons are consistent with previous studies in neighboring motor areas^52,53^. In IT neurons, we found that neurons that project both to the contralateral cortex and the striatum occupy two different sub-laminae depending on whether the neurons project to the contralateral striatum (**Extended Data Fig. 19k, l**). Therefore, the high throughput of BARseq enabled the discovery of projection patterns enriched in specific sublayers, beyond the major class-level divisions.

The large sample size of BARseq allowed us to further examine whether higher order structure in the projections (**Extended Data Fig. 19m**) is informative of the laminar locations of the neurons. Consistent with this hypothesis, the laminar positions of IT neurons were strongly dependent on projections to the secondary somatosensory cortex, but the relationship between such projections and the laminar positions of neurons were reversed in neurons with or without contralateral projections. Neurons projecting only to the ipsilateral cortex, but not the secondary somatosensory cortex (group 14), were confined to a thin layer at the bottom of L6 (**Fig. 6h; Extended Data Fig. 19n**) (this is also consistent with retrograde tracing results as shown in **Fig. 3a,b**), whereas those projecting to the ipsilateral secondary somatosensory cortex (groups 13 and 15) were mostly in L2/3. Conversely, callosal neurons projecting to the somatosensory cortex were more enriched in L6 compared to those not projecting to the secondary somatosensory cortex (**Fig. 6i; Extended Data Fig. 19o**). To test whether such dependence of laminar positions on the combinatorial projection patterns were generally true, we compared the performance of two models that predicted the laminar positions of neurons using individual projections or combinatorial projection patterns. If the higher order structure in projections did not contain additional information regarding the laminar positions of the neurons, then the two models would perform similarly in predicting projections. However, the model trained on the combinatorial projection patterns consistently outperformed the model trained on individual projections (**Fig. 6j**, p = 5×10^−31^ using t-test), indicating that the combinatorial projection patterns contained additional information regarding the laminar positions of the neurons than individual projections. Such relationships would have been difficult to find in previous attempts to identify clear projection correlates of transcriptomic neuronal subtypes using retrograde labeling of single projections^52^. Therefore, the high order structure in projection patterns predicts laminar positions of neurons.

### Classification of projection patterns revealed by complete single-cell reconstruction

Complete single cell reconstruction achieves the ground truth for morphological analysis and classification, but such datasets have been extremely difficult to obtain. We generated complete morphologies of 38 MOp neurons using systematic whole brain sparse labeling, fMOST imaging, and registration to CCFv3^54^. We augmented this dataset with 98 L5-6 MOp single neuron reconstructions from the Janelia MouseLight Project^55^, and a third set of fMOST reconstructions (n=15 cells) for a total of 151 completely reconstructed axonal arbors. For each neuron, we calculated the fraction of total axonal length within each of 314 target structures summed across hemispheres (**Fig. 7a**, matrix). We compared the distribution differences among all axonal pairs with a randomized distribution obtained by shuffling the target regions of each neuron, but preserving the number of regions invaded by each neuron (**Extended Data Fig. 20B**). The shuffled distribution is significantly narrower than the real distribution (CV_shuffled_=0.4992, half-height-width_shuffled_=16; CV_real_=0.5294, half-height-width_real_=25; p<10^−25^, Levene’s test), which disproves the ‘continuum’ hypothesis. The prominent left tail reflects the similarity between neurons within the same projection class, while the right tail corresponds to the differences between neurons from different projection classes. This analysis suggests the existence of different clusters of axon projection patterns at the single cell level. We thus performed unsupervised hierarchical clustering on these data to identify the projection classes and their relationship with the bulk anterograde tracing results (from **Fig. 4**, n=19, Extended Data **Table 1**). We identified six main clusters (C1-C6, **Fig. 7a**, dendrogram) and annotated them based on the projection classes assigned previously for Cre line tracer experiments in each cluster. The first split in the dendrogram separated the ET (C1-C3) from IT class (C4-C6). Within the ET group, the next split separates L5 ET (blue, C1-C2) from L6 CT (red, C3). Two L5 ET clusters were identified at this threshold; C1 is significantly enriched for single cells in L5 and contains all the Cre lines and wild type tracer experiments that label L5 ET cells as well as single cells projecting to many subcortical targets, including thalamus, midbrain, pons, and medulla. The smaller ET C2 has relatively weaker projections to cortical and thalamic targets. C3 is significantly enriched for single cell somas in L6 and contains the L6 CT Cre line tracer experiments (Ntsr1-Cre_GN220 and Tle4-Cre).

**Figure 7.**
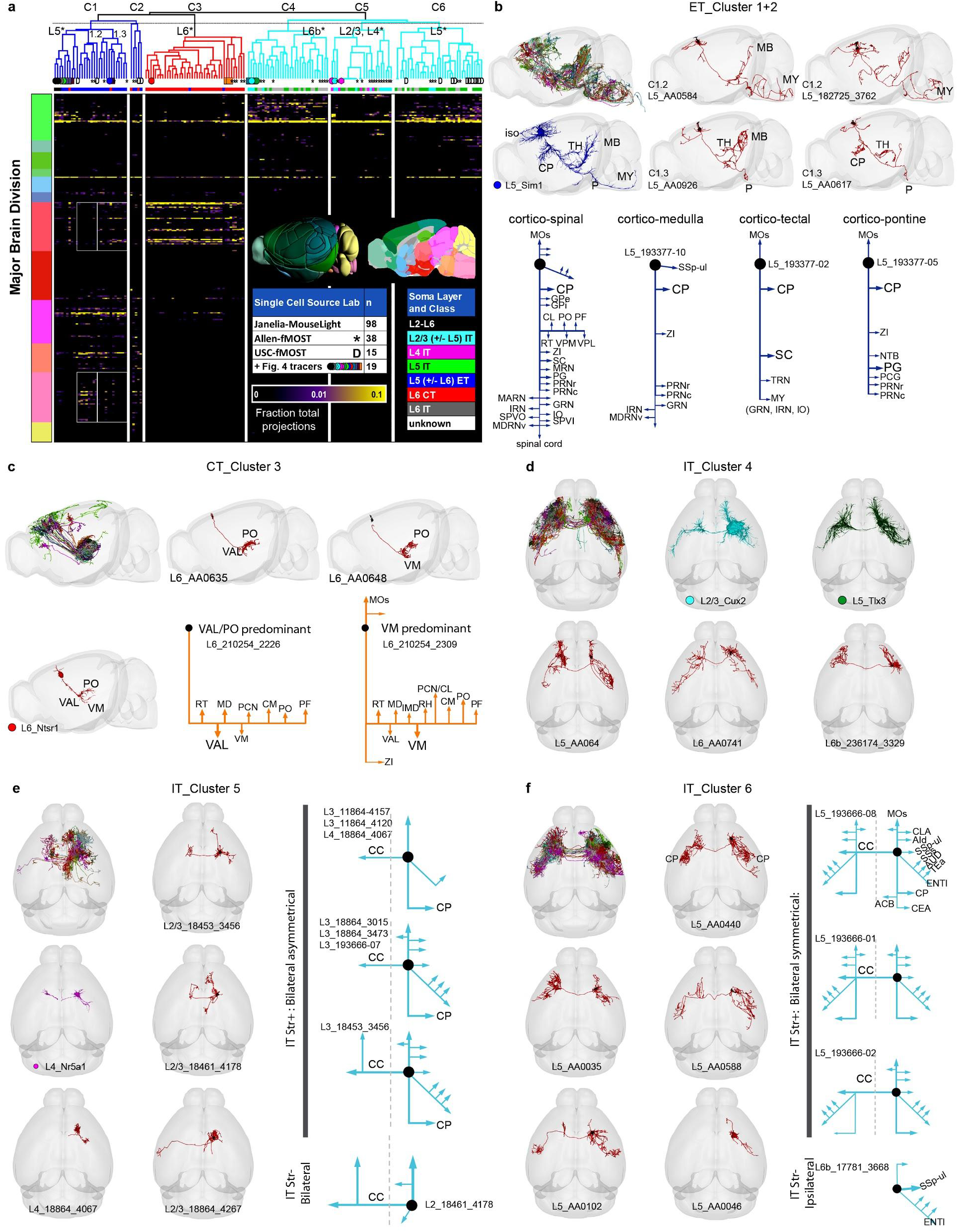
Full morphological reconstruction reveals diverse single cell projection motifs. **a**, The fraction of total axon projections from tracer and single cell reconstruction experiments to each of 314 targets across the major brain divisions. Columns show individual experiments. Rows show target regions ordered by major brain division. Hierarchical clustering and cutting the dendrogram at the level indicated with a dashed line revealed six clusters. Layer of origin for each cell or Cre line is shown under the dendrogram, colored as indicated by the “soma layer and class” inset. Cells from specific layers were significantly enriched within clusters as indicated by the * (Fisher’s exact test, p<0.05). **b**, top left, lateral view of all single cells and their axons assigned to C1 and C2 (n=26) overlaid and randomly colored reveal projections consistent with the L5 ET class. Additional lateral views show the brain-wide projection pattern from a L5 ET Cre line assigned to this cluster (Sim1-Cre, in blue) and four individual single cells (red). Several distinct L5 ET motifs were identified following visual inspection and classification of projection patterns (bottom schematics). **c**, top left, lateral view of all single cells and their axons assigned to C3 (n=38) overlaid and randomly colored reveal projections consistent with the L6 CT class. Additional lateral views show the brain-wide projection pattern from a L6 CT Cre line assigned to this cluster (Ntsr1-Cre) and two individual single L6 cells (red). Two distinct motifs were identified following visual inspection and classification of single-cell projection patterns. VAL/PO predominant neurons are distributed in L6a; while VM predominant projecting neurons are primarily distributed in L6b. **d**, top left, top view of all single cells and their axons assigned to C4 (n=29) overlaid and randomly colored reveal projections consistent with the IT class. Additional top views show brainwide projection patterns from L2/3 (Cux2-Cre) and L5 (Tlx3-Cre) IT Cre lines assigned to this cluster and three individual single cells from deep layers as indicated (red). **e**, top left, top view of all single cells and their axons assigned to C5 (n=22) overlaid and randomly colored reveal projections consistent with the IT class. Additional top views show brain-wide projection patterns from a L4 IT Cre line (Nr5a1-Cre) assigned to this cluster and four individual single cells from superficial layers as indicated (red). Several IT motifs, notably including bilateral *asymmetrical* projection patterns, were identified following visual inspection and classification of projection patterns (right schematics). **f**, top left, top view of all single cells and their axons assigned to C4 (n=36) overlaid and randomly colored reveal projections consistent with the IT class. Additional top views show five individual single cells from L5 (red). Several IT motifs, notably including bilateral relatively *symmetrical* projection patterns, were identified following visual inspection and classification of projection patterns (right schematics).

The three clusters within the IT branch (C4-C6) contain single cells from L2/3, L4, L5, and L6. C4 contains both L2/3 and L5 IT lines, Cux2-Cre, Plxnd1-CreERT2, and Tlx3-Cre_PL56. In contrast, C5 contains the two L4 Cre lines as well as the L2/3 Cre line, Sepw1-Cre. These IT clusters are further differentiated by (1) soma layer location (enriched for L6b in C4, L2/3 and L4 in C5, L5 in C6, Fisher’s exact t-test, p<0.05), (2) number of targets reached per experiment (C5 has significantly fewer than C4 and C6, one-way ANOVA and Tukey’s post-hoc test, p<0.0001 and p=0.0002, respectively), and (3) the specific targets contacted (2-way repeated measures ANOVA, p<0.0001 interaction effect of cluster × target area). For example, C4 has significantly more axonal projections compared to C6 in frontal and lateral cortical regions FRP, ORBl, ORBvl, AId, AIv, and MOs (Tukey’s post-hoc test, p <0.05), all of which receive inputs from the MOp via the frontal (rostral) pathway (**Supplementary Information**). In contrast, C6 has significantly more axonal projections than C4 to SSs and AUDd (Tukey’s post-hoc test, p <0.05), which are innervated by MOp-ul via the lateral pathway (**Supplement Information**). Finally, C5 had significantly fewer axons in CP compared to C6 (p=0.0119), and a trend in the same direction compared to C4 (p=0.0728). This difference is consistent with the retrograde labeling (Fig. 3) and Cre line tracing (**Fig. 4**) showing that L5 IT cells have more CP projections compared to L2/3.

To estimate the relative proportions of the 6 clusters we matched their respective single-cell axonal patterns against the regional patterns from PHA-L anterograde tracing across all target regions. This problem is equivalent to a set of constrained, weighted, linear equations that can be solved numerically by standard non-negative least-square (NNLS) or bounded-variable leastsquares (BVLS) optimization (Attili et al. 2020 doi.org/10.1007/s10479-020-03542-7). The results converge with minimal error (<1% residual sum of squares) on the following compositions: 47% C1, 3.5% C2, 19% C3, 4.5% C4, 1% C5, and 25% C6, corresponding to 50.5% ET, 19% CT, and 30.5% IT.

### Diverse and distinct PN axon projection motifs

As our single cell analysis revealed substantial projection pattern variability among individual neurons within the same projection class, we examined whether there were finer axon arbor motifs determining discrete neuron types. Individual CT neurons (C3) are the least diverse in their projection patterns (average Spearman R=0.66) compared to ET and IT neurons (C1/C2 R=0.39, C4-C6 R=0.45). The ET and IT correlation coefficients indicate considerable within-cluster diversity of axon branching motifs (**Fig. 7b-f**). MOp L5 ET neurons in C1/C2 project to thalamus, midbrain, pons, medulla, and spinal cord, and collateralize in cortex and striatum (**Fig. 7b**). Two L5 ET subclusters differ in their projections to either medulla (subcluster C1.2: **Fig. 7b**, top right) or thalamus (subcluster C1.3: **Fig. 7b**, bottom right), as previously reported^53^. In combination with retrograde labeling (**Fig. 3**) and BARseq data (**Fig. 6**), we further identified 4 projection categories of L5 ET cells (**Fig 7b**, bottom schematics): (1) cortico-spinal, (2) cortico-medullary, (3) cortico-tectal, and (4) cortico-pontine. We identified two categories of CT neurons in C3 with their less variable projection motifs (**Fig. 7c**): (1) those strongly projecting to ventromedial thalamic nucleus (VM), and (2) those that instead predominantly target VAL and PO. These two types of CT neurons are differentially distributed in L6b or L6a (**Fig. 3**). Both types also project significantly to several other thalamic nuclei, such as the MDl, PCN, CL and PF (**Fig. 7c**, bottom right).

IT neurons with highly variable projection motifs in C4-C6 (**Fig. 7d-f**) can be broadly divided based on whether they project to the striatum (IT Str+ and IT Str-). The IT Str+ neurons, all of which also projected to isocortex, were further differentiated based on whether they had ipsilateral or bilateral striatal connections. The ipsi IT Str+ cells (n=9) were mostly in L2/3 or L4 (8 of 9 cells) and grouped in C5 (6 of 9 cells, **Fig. 7e** right). Their cortical collaterals varied considerably in complexity, but were notably bilaterally-asymmetric. The L5 IT Str+ neurons (n=3) were clustered in C6, and displayed more or less bilaterally-symmetric projections (**Fig. 7f** right). One cell had strong bilaterally symmetric connections with several endbrain regions, including nucleus accumbens and pallidum (specifically the central amygdalar nucleus capsular part, CEAc), which has not been reported previously. The IT Str-cells are grouped in C5 and C6 (**Fig. 7e,f**, bottom schematics). Their projection patterns were either ipsilateral only or had (additionally or exclusively) contralateral connections. MOp neurons with ipsilateral-only targets connected to multiple regions, like somatosensory (SSp-ul, SSp-bfd, and SSs), auditory (AUDv) and other temporal areas (TEa, ECT, and PERI), and ENTl, or occasionally to single regions (MOs or SSp-ul). IT Str-with contralateral projections largely mirrored the projection patterns of their ipsilateral counterparts. Together, these results suggest that the substantial variation of axon projection patterns may derive, at least in part, from definable fine scale structural motifs.

To relate regional inputs and somatic layering to single-cell morphology, we compared the dendritic arbors of superficial (L2/3/4) and deep (L5) MOp pyramidal cells (**Extended Data Fig. 21a**). We reconstructed 68 neurons using two approaches: crossing MORF3 mice^56^ to Cux2-CreERT2 and Etv1-CreERT2 to label respectively superficial and deep cells (UCLA/USC); and crossing the TIGRE-MORF (Ai166) reporter line^63^ with Cux2-CreERT2, Fezf2-CreERT2, and Pvalb-Cre (AIBS). We then analyzed basal dendrites, accounting for the bulk of dendritic surface area^57^, to investigate structure beyond the expected differences in pial-reaching apical trees. Arbor morphology significantly segregated deep from superficial cells (**Extended Data Fig. 21b**): L5 neurons had larger and more complex basal trees than neurons in superficial layers (**Extended Data Fig. 21c**), as also reported for other isocortical regions in different species^64–67^. Additionally, relative to their invaded volume, superficial neurons distributed a greater proportion of their dendritic length distally from the soma compared to their deep counterparts (**Extended Data Fig. 21d**).

## Discussion

A cellular resolution anatomical framework is necessary to guide the exploration of brain organization and function. High-throughput single cell RNA sequencing provides a powerful approach to identify transcription-defined cell types and to derive a comprehensive cell census of multiple brain regions^52,58^. Furthermore, the advancement of *in situ* single-cell transcriptomics methods (e.g. MERFISH, seqFISH) will enable the generation of a molecularly defined and spatially resolved cell atlas of brain regions^44,59,68^. An essential next step is to map the morphology and, ultimately, the connectivity of these transcriptomic-defined cell types, especially PN types, within a high resolution 3D whole-brain framework^23^. Here we report the strategies, data production pipelines, and progress toward this goal. Combining multiple state-of-the-art labeling, imaging, anatomical tracing, and neuroinformatics methods, we identified the most comprehensive set of anatomically defined PN types of a mammalian cortical area (**Fig. 8a**). The distribution of these PN types reveals a much finer cortical layer definition than previously appreciated (**Figs. 3a, 8a**). We further derive a PN type-based input-output wiring diagram of the mouse MOp-ul (**Figure 8b,c**). This cell type resolution anatomic scaffold establishes a key foundation for future cellular resolution mapping of molecular information in the context of neural circuits and exploring the circuit dynamics and global network operations that underlie brain function and behavior (**Figure 8c**).

**Figure 8.**
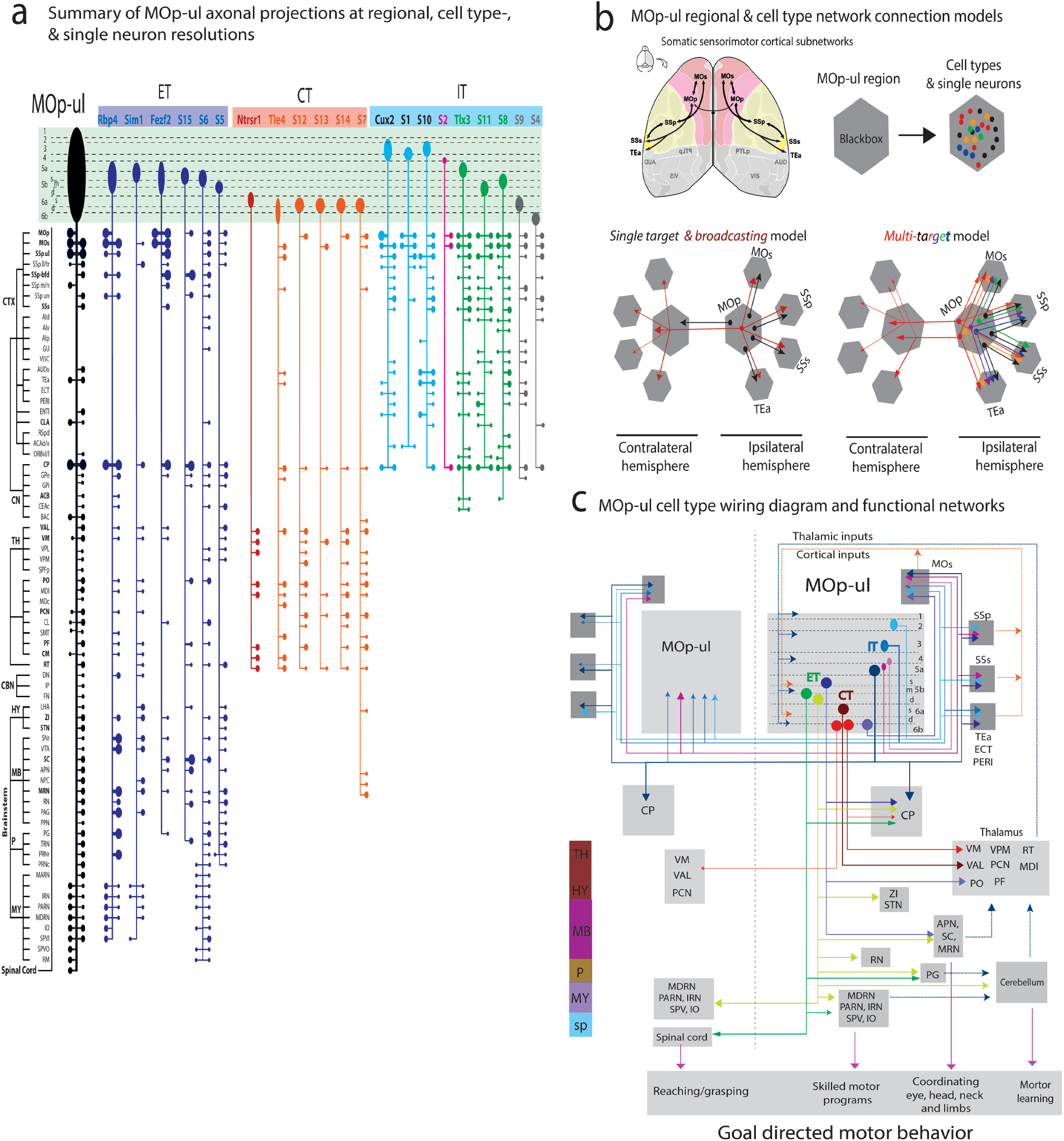
Cell type wiring diagram of the MOp. **a**, Schematic summary of projection patterns arising from major cell types (ET, CT and IT) compared with the overall projection pattern from MOp-ul (left, black). Along each vertical pathway, the output from one Cre-line tracing or single cell reconstruction (labeled “Sx”) experiment is summarized. Somatic laminar location is shown for each at the top. The horizontal bars and dots represent ipsilateral (right) and contralateral (left) axon terminals are present in the targets shown. Dot sizes indicate the relative strength of each projection. **b**, MOp-ul shares extensive connections with other cortical areas, including bilateral projections to MOs, SSp, SSs, TEa and to contralateral MOp. At regional level, MOp-ul serves as one individual network node. As shown in this study, the MOp-ul contains more than 15 types of IT neurons with distinctive connections, which drastically increase complexity of network models. The two network schematics show three hypothetic models describing IT cell type-specific cortical connections; bottom left side (1), a *single target* connection model (black), in which a cell type innervates only one cortical target (e.g., MOp-ul-->MOs) and (2) a *broadcasting model*, in which a cell type innervates many cortical targets (7-11 targets) of MOp (red); bottom right side (3) *a multi-target model*, in which a cell type projects most commonly to 2-4 targets with different combinations. Note that in this model (and in the case of multiple models coexisting), one cortical target receives inputs from multiple types of MOp-IT neurons. **c**, A summary wiring diagram of MOp-ul cell-types and predicted functional roles. A subset of cortical and striatal projection patterns are shown from the diverse MOp IT cell types (n=6 representing cells in L2 – L6b). Three types of MOp-ul CT neurons are shown based on the specific combination of thalamic targets and sublayer soma location. Summary projection patterns from three types of ET neurons are also illustrated here. Each type projects predominantly to different subcortical targets involved in distinct motor functions. (1) c*ortico-spinal projection neurons* to the cervical spinal cord in controlling goal-directed upper limb motor activities, such as reaching and grasping^69^; (2) projections to the reticular formation (e.g. MDRN) from *cortico-medullar* neurons are implicated in task-specific aspects of skilled motor programs^70^; and (3) *cortico-tectal* projection to the SC is implicated in coordinating movements of eye, head, neck, and forelimbs during navigation and goal-oriented behaviors (such as escaping or appetite behavior)^71^. All L5 ET neurons and IT neurons generate projections to the striatum; and ‘IT/PT balance’ were implicated in a range of neuropsychiatric, neurodevelopmental and neurological disorders^41^.

Classic studies of brain organization have defined macroscale network nodes at the level of gray matter regions, such as “areas”, “layers”, and “nuclei”, based largely on cyto-and histoarchitecture. However, precise spatial boundaries of many such divisions are often unclear or inconsistent (e.g. the MOp-ul), casting inherent limitations on the reliability of such macroscale divisions and connectomes. In contrast, cell type resolution is a key level to define mesoscale network nodes that should clarify historical ambiguities and controversies. In particular, a foundational concept of cortical architecture is the notion of a six-layered laminar organization, based on then available histological, molecular, and developmental evidence. While this concept has guided cortical research for several decades, there is increasing evidence that defies a simple and strict six-layer organization. Here, by resolving a comprehensive set of PN types and their spatial distribution, we reveal a substantially finer scale laminar organization. Consistent with our findings, a recent in situ single-cell transcriptomics analysis of molecularly defined cell types also revealed a similar finer sublaminar organization of cell types^44^, further supporting our refinement of a largely cytoarchitonics-based six-layer cortical organization to eleven sublayers. In other regions of the nervous system where cell types are better recognized, cellular resolution analyses have delineated ten layers in the retina^60^ and dozens of layers and spatial domains in the spinal cord^61^. More generally, it is likely that the distribution pattern and connectivity of neuronal cell types, increasingly revealed by emerging datasets, may substantially clarify and refine classic macroscale descriptions of brain regions and their subdivisions (e.g.,layers, cores and shells).

A rigorous definition of PN types, and neuron types in general, is by no means settled. Single cell RNAseq studies have identified approximately 50 transcription-defined glutamatergic neuron types in several cortical areas, including the MOp, most of which are PNs^52^,^58^. Here, based largely on projection patterns and molecular markers, we define 25 MOp-ul PN types. A major goal is to achieve a multi-modal correspondence, with appropriate granularity, to define cell types. From our anatomical data modalities using, Cre driver line-based anterograde, retrograde and trans-synaptic tracing, we identified over two dozen PN subpopulations. However, in our opinion, large scale complete single cell reconstruction provides the ultimate desired resolution to achieve ground truth discovery and an internally consistent classification of anatomic cell types. With the availability of reliable driver lines targeting major PN classes, a saturation screen of single neuron morphology within each class is now feasible, especially with continued technology development^56, 63^. Despite exclusively focusing on PNs, the number of neuron types identified here exceeds earlier estimates^7^ by 5-fold. Although differences among brain regions are expected, our results underscore the enormous diversity of long-range glutamatergic neuron types besides the more readily recognized diversity of cortical GABAergic interneurons^62^.

Knowledge of evolutionary conservation and divergence of brain structures often yields insights into core organizational principles. Previous cross-species comparisons of mammalian brains have largely been carried out at the level of gray matter regions, such as areas and layers in the cortex, leaving many open questions on what is and is not conserved. The joint molecular and anatomic identification of PNs will provide a high resolution and robust metric for cross-species translation. Although the primate cortex has more functionally distinct areas and potentially orders of magnitude larger cortical networks than those in rodents, a PN type resolution analysis may reveal truly conserved core subnetworks and novel species innovations. The MOp may provide a good starting point for such comparative studies, given the clearly recognizable conservation and divergence of forelimb structures and motor behaviors from rodents to humans. Charting a PN type-based circuit diagram of a small region of the mouse brain points out several key future studies. First, the strategy and pipeline shaped by our study of mouse MOp-ul will apply to other cortical and brain regions, including the whole mouse brain. Second, large-scale single neuron complete reconstructions, especially when linked to class-defining molecular markers, is both feasible and crucial to achieve a ground truth definition of PN types. A major challenge still to overcome at scale is the systematic integration of molecular, spatial, and projection information at the level of cell types and single cells. Retrograde labeling combined with in situ single cell transcriptomics provides one approach^44^, but other new methods are needed. Furthermore, a high resolution synaptic wiring diagram requires accurate identification of pre- and post-synaptic neurons, ideally at single cell resolution. This will require new technologies or advances in scaling up electron microscopy for the mouse brain^72^. Such cellular resolution structure framework is a crucial step toward a “Google Map of the brain”, where the seamless “zooming in-and-out” exploration across different levels of brain organization can be achieved by tracking the key information processing scaffold upon which multi-modal biological properties are organized and through which physiological computations are played out.

## Acknowledgment

B.Z., H.H., H.W.D., I.B., L.G., L.G., L.K., M.S.B., M.Z., N.N.F. thank Kaelan Cotter, Luis Gacia, Darrick Lo, Tyler Boesen, Chunru Cao, Marlene Becerra, Marina Fayzullina, Christine Mun for their technical and informatics support. Their work was supported by NIH U01MH114829 (to H.W.D., G.A.A. and B.L.), R01MH094360 (H.W.D.), U19MH114821 (J.H./P.A.), and U19MH114831 (J.E./E.C.).

A.B., A.N., C.E., J.M., J.P., P.O., R.D., R.M.C., R.P. and X.Q. were supported in part by NIH U01 MH114824 to P.O.

D.W.W., S.M.A., and G.A.A. gratefully acknowledge the assistance of Tiago Ferreira in accessing the API for batch downloading the Janelia MouseLight neuron dataset and for providing constructive feedback on the corresponding analysis. Their work was supported by NIH U01MH114829 (to H.W.D., G.A.A. and B.L.), R01NS39600 (to G.A.A.), and R01NS86082 (to G.A.A. and Dan Cox).

A.C., E.F., F.D., H.Z., J.A.H., K.E.H, L.M, M.N., M.H., P.A.G., P.L., P.R.N., Q.W., S.N., S.Y., W.W., and Y.W. are grateful to the Transgenic Colony Management, Neurosurgery & Behavior, Lab Animal Services, Molecular Genetics, Imaging, and Histology teams at the Allen Institute for technical support. In particular, they thank Vonn Wright, Medea McGraw, Lydia Potekhina, Leonard Kuan, and Ali Williford from these teams. Their work was supported by the Allen Institute for Brain Science and by NIH grants R01EY023173, U01MH105982, and U19MH114830 to H.Z. Those authors thank the Allen Institute founder, Paul G. Allen, for his vision, encouragement, and support.

Y.C.S, A.M.Z, and X.C. acknowledge members of the MAPseq core facility, including Huiqing Zhan and Yan Li, for facilitating BARseq data production, and Wiktor Wadolowski for technical support. Their work was supported by NIH 5R01NS073129, 5R01DA036913, RF1MH114132, and U01MH109113 to A.M.Z, R01MH113005 and R01LM012736 to J.G., and U19MH114821 to A.M.Z. and J.G.; the Brain Research Foundation (BRF-SIA-2014-03 to A.M.Z.), IARPA MICrONS (D16PC0008 to A.M.Z.), Paul Allen Distinguished Investigator Award (to A.M.Z.), Simons Foundation (350789 to X.C.), Chan Zuckerberg Initiative (2017-0530 ZADOR/ALLEN INST(SVCF) SUB to A.M.Z), and Robert Lourie (to A.M.Z.). Their work was additionally supported by the Assistant Secretary of Defense for Health Affairs endorsed by the Department of Defense through the FY18 PRMRP Discovery Award Program W81XWH1910083 to X.C. Opinions, interpretations, conclusions, and recommendations are those of those author and are not necessarily endorsed by the U.S. Army. In conducting research using animals, the investigator adheres to the laws of the United States and regulations of the Department of Agriculture.

B.Z., J.H., H.W.T, and L.I.Z. were also supported by NIH R01DC008983, RF1MH114112, U01MH116990, and EY019049.

J.G. was supported by NIH R01NS096720.

J.T.H., K.K., K.S.M., W.G., X.A., and Z.J.H. were supported in part by NIH 5U19MH114821-03 to Z.J.H.

P.M. was supported by NIH EB022899, MH114824, MH114821, and NS107466, the Mather Foundation, and a Crick-Clay Professorship.

X.W.Y. and H.D. were supported by NIH BRAIN Initiative MH106008; X.W.Y., H.D., M.Z., and N.F. were supported by NIH BRAIN Initiative MH117079.

Y.K. was supported by NIH R01MH116176.

H.P., L.L., P.X., L.D., and Y.W. were supported by an Open Science initiative at Southeast University.

## METHODS

### Dong Lab

#### Data Production

##### Mouse Connectome Project methodology: Multiple fluorescent anterograde, retrograde, and viral tracing

Anatomical tracer data was generated as part of the Mouse Connectome Project (MCP). MCP experimental methods and online publication procedures have been described previously^1,2^. To systematically map the input and output neuronal connectivity of the MOp-ul, we used a multi-fluorescent tracing strategy with a combination of classic tract-tracing and viral tracing methods. First, we used a double co-injection approach that injects two different tracer cocktails each containing one anterograde and one retrograde tracer to simultaneously visualize two sets of input/output connectivity. Then, we used a multiple retrograde tracing method to simultaneously inject different fluorophore-conjugated retrograde tracers into different projection targets (up to 4) of the MOp in the brain and spinal cord to retrogradely label projection neurons in the MOp. Finally, we used a triple anterograde tracing approach with individual injections of three different anterograde tracers into three different brain targets.

##### Animal subjects

All MCP tract-tracing experiments were performed using 8-week old male C57BL/6J mice (Jackson Laboratories). For anterograde trans-synaptic tracing experiments and AAVretro injections in spinal cord, Ai14 tdTomato Cre-reporter mice were used (Jackson Laboratories, stock #007914, aged 2-3 months old). Mice had ad libitum access to food and water and were group-housed within a temperature- (21-22°C), humidity- (51%), and light- (12hr: 12hr light/dark cycle) controlled room within the Zilkha Neurogenetic Institute vivarium. All experiments were performed according to the regulatory standards set by the National Institutes of Health Guide for the Care and Use of Laboratory Animals and by the institutional guidelines described by the USC Institutional Animal Care and Use Committee.

##### Tracer injection experiments

The MCP uses a variety of combinations of anterograde and retrograde tracers to simultaneously visualize multiple anatomical pathways within the same Nissl-stained mouse brain. Triple anterograde tracing experiments involved three separate injections of 2.5% *Phaseolus vulgaris* leucoagglutinin (PHAL; Vector Laboratories, Cat# L-1110, RRID:AB_2336656), and adeno-associated viruses encoding enhanced green fluorescent protein (AAV-GFP; AAV2/1.hSynapsin.EGFP.WPRE.bGH; Penn Vector Core) and tdTomato (AAV1.CAG.tdtomato.WPRE.SV40; Penn Vector Core). Retrograde tracers included cholera toxin subunit B conjugates 647, 555 and 488 (CTb; AlexaFluor conjugates, 0.25%; Invitrogen), Fluorogold (FG; 1%; Fluorochrome, LLC), and AAVretro-EF1a-Cre (AAV-retro-Cre; Viral Vector Core; Salk Institute for Biological Studies). To provide further details on specific connectivity patterns, we also performed quadruple retrograde tracer, and rabies/PHAL experiments. Quadruple retrograde tracer experiments involved four different injections sites receiving a unique injection of either 0.25% CTb-647, CTb-555 CTb-488, 1% FG, or AAV-retro-Cre. Retrograde tracing from the spinal cord (Fig. 2d; Extended Data Fig. 2.3) was performed with AAVretro-hSyn-GFP-WPRE (Addgene, Cat# 50465) and AAVretro-hSyn-Cre-WPRE (Addgene, Cat# 105553). To further establish synaptic connectivity in downstream targets of MOp-ul (Extended Data Fig.4.2), AAV-hSyn-mRuby2-sypEGFP (Custom design, Laboratory of Byunkook Lim) was used to label axons-of-passage with mRuby2 (red) and presynaptic puncta with EGFP (green). Patterns of synaptic innervation were further demonstrated using injections of self-complementary (sc) AAV1-hSyn-Cre (Vigene Biosciences; 2.8 x 10^13^ GC/ml), which is capable of anterograde transneuronal spread to post-synaptic targets^3,4^. To reveal mono-synaptic inputs to a projection defined neuronal populations (Figure 5; Extended data Fig. 17), we used a modified TRIO (tracing the relationship between input and output) strategy (Schwarz et al., 2015). In brief, AAVretro-Cre was injected into a MOp downstream projection target (i.e., caudoputamen) and Cre-dependent, TVA- and RG-expressing helper virus (AAV8-hSyn-FLEX-TVA-P2A-GFP-2A-oG) and mCherry-expressing G-deleted rabies virus (produced by Laboratory of Ian Wickersham at MIT) are injected into the MOp to label the MOp projection neurons population (1^st^-order) and their brain-wide monosynaptic inputs (2^nd^-order).

All cases used in this study are listed in Extended Data Tables 2 and 3. No statistical methods were used to pre-determine sample sizes, but our sample sizes are similar to those reported in previous publications. In most cases, anterograde tracing results are cross validated by retrograde labeling injections at anterograde fiber terminal fields and vice versa. Data generated is published online as part of the Mouse Connectome Project (www.MouseConnectome.org).

##### Stereotaxic surgeries

On the day of the experiment, mice were deeply anesthetized and mounted into a Kopf stereotaxic apparatus where they were maintained under isofluorane gas anesthesia (Datex-Ohmeda vaporizer). For triple anterograde injection experiments, PHAL was iontophoretically delivered via glass micropipettes (inner tip diameter 24-32μm) using alternating 7sec on/off pulsed positive electrical current (Stoelting Co. current source) for 10min, and AAVs were delivered via the same method for 2 min (inner tip diameter 8-12μm). For anterograde/retrograde coinjection experiments, tracer cocktails were iontophoretically delivered via glass micropipettes (inner tip diameter 28-32μm) using alternating 7sec on/off pulsed positive electrical current (Stoelting Co. current source) for 10 min (PHAL/CTB-647). For quadruple retrograde tracing experiments, 50nl of retrograde tracers were individually pressure-injected via glass micropipettes at a rate of 10nl/min (Drummond Nanoject III). Pressure injections were also similarly performed for anterograde transsynaptic experiments using scAAV1-hSyn-Cre (100 nL), labeling of presynaptic boutons with AAV-hSyn-mRuby2-sypEGFP (50 nL), and spinal injections of AAVretro-hSyn-GFP-WPRE and AAVretro-hSyn-Cre-WPRE (cervical and lumbar, respectively, 200 nL each; see Zingg et al., 2020 for detailed procedure). For TRIO experiments, AAVretro-hSyn-Cre-WPRE was injected into one MOp-ul downstream projection target (i.e., caudoputamen). And, Cre-dependent, TVA- and RG-expressing helper virus (AAV8-hSyn-FLEX-TVA-P2A-GFP-2A-oG) was injected into the MOp-ul. Animals were then survival for 2 weeks before receiving injections of mCherry-expressing G-deleted rabies virus in the MOp-ul. All injections were placed in the right hemisphere. Injection site coordinates for each surgery case are on the Mouse Connectome Project iConnectome viewer (www.MouseConnectome.org).

##### Histology and immunohistochemical processing

After 1-3 weeks post-surgery, each mouse was deeply anesthetized with an overdose of Euthasol (pentobarbital) and trans-cardially perfused with 50ml of 0.9% saline solution followed by 50ml of 4% paraformaldehyde (PFA, pH 9.5). Following extraction, brain tissue was postfixed in 4% PFA for 24-48hr at 4°C. Fixed brains were embedded in 3% Type I-B agarose (Sigma-Aldrich) and sliced into four series of 50μm thick coronal sections using a Compresstome (VF-700, Precisionary Instruments, Greenville, NC) and stored in cryopreservant at −20°C. For double coinjection experiments, one series of tissue sections was processed for immunofluorescent tracer localization. For PHAL or AAVretro-EF1a-Cre immunostaining, sections were placed in a blocking solution containing normal donkey serum (Vector Laboratories) and Triton X-100 (VWR) for 1 hr. After rinsing in buffer, sections were incubated in PHAL primary antiserum (1:1000 rabbit anti-PHAL antibody (Vector Laboratories Cat# AS-2300, RRID:AB_2313686)) or AAVretro-EF1a-Cre primary antiserum (1:1000 mouse-anti-Cre) mixed with blocking solution for 48-72 hours at 4°C. Sections were then rinsed again in buffer solution and then immersed in secondary antibody solution (blocking solution and 1:500 donkey anti-mouse IgG conjugated with Alexa Fluor 488 (Thermo Fisher Scientific Cat# A-21202, RRID:AB_141607), or 1:500 donkey anti-rabbit conjugated with CY3 (Jackson ImmunoResearch Labs Cat# 715-165-151, RRID:AB_2315777) for 3 hrs. Finally, all sections were stained with Neurotrace 435/455 (Thermo Fisher Cat# N21479) for 2-3 hours to visualize cytoarchitecture. After processing, sections were mounted onto microscope slides and cover slipped using 65% glycerol.

#### Data Collection

##### Imaging and post-acquisition processing

Complete tissue sections were scanned using a 10X objective lens on an Olympus VS120 slide scanning microscope. Each tracer was visualized using appropriately matched fluorescent filters and whole tissue section images were stitched from tiled scanning into VSI image files. For online publication, raw images are corrected for left-right orientation and matched to the nearest Allen Reference Atlas (ARA) levels. An informatics workflow was specifically designed to reliably warp, reconstruct, annotate and analyze the labeled pathways in a high-throughput fashion through our in-house image processing software Connection Lens. Threshold parameters were individually adjusted for each case and tracer, resulting in binary image output files for quantitative analysis. Adobe Photoshop was used to correct conspicuous artifacts in the threshold output files that would have spuriously affected the analysis. A separate copy of the atlas-registered TIFF image files was brightness/contrast adjusted to maximize labeling visibility and images were then converted to JPEG file format for online publication in the Mouse Connectome Project iConnectome viewer (www.MouseConnectome.org).

##### Assessment of injection sites

All injection cases included in this work are, in our judgment, prototypical representatives of each brain area. We have previously demonstrated our targeting accuracy with respect to injection placement, our attention to injection location, and the fidelity of labeling patterns derived from injections to the same location (see Supplementary Methods in Hintiryan et al., 2016 for details). In the current report, we also demonstrate our injection placement accuracy and the consistent labeling resulting from injections placed in the same areas.

#### Data Analysis

The registered, thresholded image files were quantified using our in-house software Connection Lens, where each section was matched and warped to its corresponding atlas level of the Allen Reference Atlas (ARA) and the labeling was segmented. The pixels of thresholded tracer labeling were quantified in each nucleus and cortical region defined in the atlas level to which the image was registered. Results were output in a spreadsheet for statistical analysis and matrix visualization.

#### Data Presentation

Atlas registered TIFF image files were converted into JPEG2000 image format, while thresholded images were aggregated into SVG images. All fluorescently labeled connectivity data are presented through the iConnectome viewer, the iConnectome Map Viewer, and published to the Data Repository Dashboard page, www.MouseConnectome.org. Quantified cell count files and projection matrix also accessible from www.MouseConnectome.org.

#### Informatics Tools and Code availability

iConnectome Viewer and iConnectome Map Viewer are accessible from the data repository page hosted on www.MouseConnectome.org. Publicly accessible Data Repository Dashboard Page available at http://www.mouseconnectome.org/Dinoskin/page/dashboard. Public code repositories stored in GitHub (github.com).

#### Cell type atlasing, distribution and morphology analysis by the Osten Lab at CSHL Cre-reported lines

All animal procedures were performed under the Cold Spring Harbor Laboratory Institutional Animal Care and Use Committee (IACUC) approval. Animals were given food and water ad libitum and housed under constant temperature and light conditions (12 hr cycle lights ON: 0600, lights OFF: 1800). For “knock-in” animals, we crossed Cre drivers with reporter mice (CAG-LoxP-STOP-LoxP-H2B-GFP) as previously described.

#### Brain samples preparation and imaging of cell-type distributions

Cre-reported transgenic mice were anesthetized with a mix of ketamine/xylazine and perfused transcardially with isotonic saline followed by 4% paraformaldehyde (PF A) in 0.1M phosphate buffer (PB, pH 7.4). After extraction, brains were post-fixed overnight at 4C in the same fixative solution, and stored at 0.05M PB until imaging.

Before imaging, brains were embedded and cross-linked with oxidized 4% agarose as previously described ^5–7^. Whole brain imaging of Cre-reported lines was achieved using the automated whole-mount microscopy STPT. The entire brain was coronally imaged at an X,Y resolution of 1μm and Z-spacing of 50μm^5,6,8^.

#### Whole mount Neurotrace staining

Whole brain Neurotrace staining was performed with a modification of iDISCO+ protocol^9^ (Muñoz-Castañeda & Osten; manuscript in preparation).

#### STPT cell counting

Automatic cell counting in MOp-ul was done as previously described^5,6,8^. A convolutional neural network (CN) was trained using H2B-GFP nuclear signaling. First, we develop an unsupervised detection algorithm for cell detection based in structure tensor and connected components analyses. Results were used to automatically generate 270 random segmented image tiles from 3 different datasets (~1350 cells) which were posteriorly used as the ground truth (Muñoz-Castañeda & Osten; manuscript in preparation).

#### ARA Nissl registration

2D ARA Nissl slices were registered onto the Common Coordinate Framework (CCF) reference brain. Briefly, initially ARA 2D slices were pre-aligned to a subset of CCF slices with 100um spacing, giving a total of 132 slices, same than ARA (custom python script). After 2D alignment, a 3D affine transformation was apply followed by a 3D B-spline transformation (see *Image registration* below; **Extended data Fig. 2.2**)

#### Image registration

Whole brain 3D datasets were registered to the CCF reference brain. Briefly, first 3D affine transformation was calculated and followed by a 3D B-spline transformation. Similarity was computed using Advanced Mattes mutual Information metric by *Elastix* registration toolbox^10^.

2D datasets pre-registered to ARA were initially aligned using the output transformations from original ARA Nissl 2D alignment. Non-pre-registered 2D datasets were initially pre-align (see *ARA Nissl registration* above for description; **Supplementary Video 1**).

#### Anatomical features enhancing

In order to improve whole brain registration, both CCF and whole brain datasets were pre-processed to enhance intrinsic anatomical features (see below). Main anatomical features in the reference brain were initially enhanced (custom Matlab scripts). Finally, a Sobel operator was applied in order to reduce noise and computational cost during image registration process (custom Python scripts). Brain image datasets were enhanced following the same process (Muñoz-Castañeda & Osten; manuscript in preparation).

#### High resolution image registration transformation

After image registration, output transformations were used to generate high resolution registered datasets (custom Matlab scripts). Briefly, we automatically generated the displacement field of the initial registration, which were used to compute the high-resolution registration tranformations (**Supplementary Video 2**; Muñoz-Castañeda & Osten; manuscript in preparation).

#### Cloud-based visualization and delineation with Neuroglancer

Brains registered at high-resolution were converted and stored in ‘precomputed’ format in Google Cloud Platform using Cloud-Volume (https://github.com/seung-lab/cloud-volume). Cloud-based visualization was done using Neurodata’s fork (https://viz.neurodata.io/) of Google Inc Neuroglancer WebGL-based viewer (https://github.com/google/neuroglancer)^11,12^. Cloudbased delineation of MOp-ul was done using Neuroglancer’s annotation tool on the top of high-resolution registered datasets (**Supplementary Video 1**).

#### MOp-ul 3D rendering

MOp-ul annotations were exported from Neuroglancer and converted to binary image files using custom scrcipts (Python). Cortical layers were delineated based on Cell types distribution. For depth distribution analysis, MOp-ul was divided in 50μm thickness bins equally spaced between pia surface and corpus callosum. Finally, MOp-ul images were 3D rendered using ParaView software^13^.

#### Data Production

##### Animal subjects

All animal procedures were approved by Cold Spring Harbor Laboratory’s Institutional Animal Care and Use Committee (IUCAC) protocol number 18-15-12-09-12. Animals were given food and water ad libitum and housed under constant temperature and light conditions (12 hr cycle lights ON: 0600, lights OFF: 1800).

##### Tracer injection experiments

Adeno-associated viruses (AAV) serotype 9 were delivered by tail vein injection. Briefly, 30ul of rAAV9/CAG-Flex-GFP (UNC Chapel Hill, 3.7×10^12 viral particles/ml) virus was injected in the tail vein of eight-week old Emx-cre+/- mice (JAX - 005628). Five weeks after the virus injection, the mice were transcardially perfused with cold saline (0.9% NaCl) followed by formaldehyde (4% w/v of PFA). The brains were dissected out and post-fixed in formaldehyde solution overnight. The brains were then washed and stored in 0.05M PB solution until clearing.

#### Data Collection

##### Brain sample preparation and imaging

The brains were cleared using 0M Cubic solution and then imaged using oblique light sheet tomography^14^. The raw tiff images were stitched using the BigStitcher plugin in ImageJ^15^. To perform soma detections and automated reconstruction, the raw images were segmented using a topological preserving 3D U-Net convolutional neural network architecture^16^.

#### Data Analysis

The soma coordinates of all the neurons were obtained using a multi-step process that involved an initial clustering process based on thresholding of the raw volume counts followed by a binary classification of the clustered pixels using a convolutional neural network that used two channels (cropped volume of the raw channel as first channel and segmented as the second) as the input.

An isotropic 25um tiff image was generated using the Fuse dataset option in the bigStitcher plugin. This isotropic image in the oblique orientation was then transformed to obtain the coronal orientation. The coronal image was then registered onto the CCF space using elastix and transformix. Soma positions that were located within the MOpul were then identified and used for reconstruction.

To obtain a highly accurate reconstruction, stitching parameters of a local 2×2 raw volume surrounding the soma were re-calculated again using the bigStitcher plugin. Fused images of this 2×2 raw volumes were generated using the bigStitcher plugin. Fused images of the segmentation volumes were also generated using the bigStitcher plugin from the same stitching parameters. The soma co-ordinates were then transformed to the new co-ordinates obtained from the local stitching transformations. These new soma co-ordinates were then used as the seed points to reconstruct the neurons.

While the all the neurites were segmented, they were often discontinuous in the segmentations. An algorithm, that used the soma co-ordinate in the raw image as the seed point, while taking into account the various parameters such as intensity of the corresponding pixel in both raw and the segmentations, intensity gradient along the raw image volume, distance to propagate if there are no segmentations, was used to grow/connect the discontinuous branches in the segmentation volume. Furthermore, a pruning algorithm that used the minimum threshold length and radius as the variables was used to prune branches. G-Cut algorithm was used to remove bridges between multiple neurons^17^. The swcs were then normalized and registered for further dendritic analysis. All animal procedures were approved by Cold Spring Harbor Laboratory’s Institutional Animal Care and Use Committee (IUCAC) protocol number 18-15-12-09-12.

#### Data Production at the Allen Institute: Viral Tracer Experiments

##### Animal subjects

Male and female transgenic mice at an average age of P56 were utilized for all experiments (viral tracer and single neuron reconstructions). Mice had ad libitum access to food and water and were group-housed (3-5 per cage) within temperature (21-22°C), humidity (40% Rh), and light (12hr: 12hr light/dark cycle) controlled rooms within the Allen Institute vivarium. Experiments involving mice were approved by the Institutional Animal Care and Use Committees of the Allen Institute for Brain Science in accordance with NIH guidelines. Wildtype adult male C57BL/6J mice (stock 00064, The Jackson Laboratory) were used for panneuronal anterograde viral tracing experiments. Cre lines were used for anterograde and retrograde viral tracing experiments and single neuron reconstructions. Cre lines were originally derived on various backgrounds, but most were crossed to C57BL/6J mice > 10 generations and maintained as heterozygous lines. Transgene expression patterns in many Cre driver lines used in this study were previously described^18,19^ and are available through the Transgenic Characterization data portal (http://connectivity.brain-map.org/transgenic).

##### Viral tracers

Whole brain axonal projections from MOp-ul were labeled with anterograde rAAV using the previously established Allen Mouse Brain Connectivity Atlas pipeline. Experimental methods and procedures have been described previously^18,19^. In brief, a pan-neuronal AAV expressing EGFP (rAAV2/1.hSynapsin.EGFP.WPRE.bGH, Penn Vector Core, AV-1-PV1696, Addgene ID 105539) was used for stereotaxic injections into wildtype C57BL/6J mice. To label genetically-defined populations of neurons, we used a Cre-dependent AAV vector that robustly expresses EGFP within the cytoplasm of Cre-expressing infected neurons (AAV2/1.pCAG.FLEX.EGFP.WPRE.bGH, Penn Vector Core, AV-1-ALL854, Addgene ID 51502).

For retrograde mono-synaptic whole brain tracing of inputs to Cre-defined cell types in MOp-ul, we used a dual virus strategy^20^. A Cre-dependent rAAV helper virus co-expressing TVA receptor, rabies glycoprotein (G), and tdTomato in the cytoplasm of Cre-expressing infected neurons (AAV1-Syn-DIO-TVA66T-dTom-N2cG) was stereotaxically injected, followed 21 +/- 3 days layer by another injection in the same location of a G-deleted, ASLV type A (EnvA) pseudotyped rabies virus expressing a nuclear GFP reporter (RV.CVS-N2c(deltaG)- H2bEGFP, Allen Institute^21^ (manuscript in progress).

##### Stereotaxic surgeries

On the day of experiment, mice were anesthetized with 5% isoflurane and placed into a stereotaxic frame (Model# 1900, Kopf, Tujunga, CA). The isoflurane level was maintained at 1-5% throughout the surgery. For anterograde viral tracer experiments and for helper virus delivery for retrograde trans-synaptic tracing, rAAV was delivered by iontophoresis with current settings of 3 or 5 μA using 7 sec on/off cycles for 5 min total, using glass pipettes (inner tip diameters of 10–20 μm). For retrograde trans-synaptic tracing, rabies virus was delivered by pressure injection (Nanoject II Variable Volume Automatic Injector, VWR) in 23 nl increments over a 3 min and 10 sec interval to a final volume of 500 nl. All injections were placed in the right hemisphere. MOp-ul was targeted using stereotaxic coordinates anterior/posterior (AP) 0.62 mm from Bregma, medial/lateral (ML) 1.5 mm from the midline at Bregma, and dorsal/ventral (DV) 0.3 or 0.6 mm from the pial surface of the brain.

##### Histology and immunohistochemical processing

Anterograde tracing brains were collected 3 weeks from the rAAV injection date. Retrograde trans-synaptic input mapping brains were collected 9 days from the rabies virus injection date. All mice were deeply anesthetized with 5% isoflurane and intracardially perfused with 10 ml of 0.9% saline solution followed by 50 ml of 4% paraformaldehyde (PFA) at a flow rate of 9 ml/min. Brains were rapidly dissected and post-fixed in 4% PFA at room temperature for 3-6 hours and overnight at 4°C. Brains were then rinsed briefly with Phosphate Buffered Saline (PBS) and stored in PBS with 0.02% sodium azide until tissue preparation for serial two photon imaging (see Imaging and post-acquisition processing below).

For rabies starter cell imaging and counting, sections were collected following serial two-photon imaging, mounted on gelatin coated glass slides and processed for immunofluorescence using an automated slide stainer (Biocare, IntelliPATH FLX). Slides were placed in a blocking solution of Image iT FX Signal Enhancer (Thermo Fisher Scientific Cat# I3693) for 45 minutes, followed by a blocking solution containing 1% normal goat serum (Vector Laboratories Cat#S1000) and 1% Triton X (VWR) for 1 hour. Sections were then incubated in dsRed primary antibody solution (1% goat serum, 1% Triton X, 1:2000 rabbit anti-dsRed antibody (Rockland Cat# 600-401-379, RRID:AB_2209751) for 1.5 hours at room temperature. Slides were rinsed in 0.1% Triton X wash solution and then incubated in secondary antibody solution (1% goat serum, 1% Triton X, and 1:500 goat anti-rabbit conjugated with Alexa Fluor 594 (Thermo Fisher Scientific Cat# A-11037, RRID:AB_2534095) for 2 hours. Finally, all sections were stained with 5 μM Dapi (Thermo Fisher Scientific D1306) and coverslipped using Fluoromount G (Southern Biotech Cat# 0100-01B).

##### Imaging and post-acquisition processing

###### Whole brain serial two-photon tomography (STPT)

STPT imaging procedures have been previously described^8,18^ (TissueCyte 1000, TissueVision Inc. Somerville, MA). In brief, following AAV tracer injections, brains were imaged at high x-y resolution (0.35 μm x 0.35 μm) every 100 μm along the rostrocaudal z-axis. Images of rabies-tracer labeled nuclei were also collected every 100 μm, but were imaged at 0.875 μm x 0.875 μm x-y resolution. Images underwent QC and manual annotation of injection sites, followed by signal detection and registration to the Allen Mouse Brain Common Coordinate Framework, version 3 (CCFv3) through our informatics data pipeline^22,23^ (IDP). The IDP manages the processing and organization of the images and quantified data for downstream analyses. The two key algorithms in the IDP are signal detection and image registration. For segmentation, high-threshold edge information was combined with spatial distance-conditioned low-threshold edge results to form candidate signal object sets. The candidate objects were then filtered based on their morphological attributes such as length and area using connected component labeling. In addition, high intensity pixels near the detected objects were included into the signal pixel set. Detected objects near hyper-intense artifacts occurring in multiple channels were removed. The output is a full resolution mask that classifies each pixel as either signal or background. An isotropic 3D summary of each brain is constructed by dividing each image series into 10 μm × 10 μm × 10 μm grid voxels. Total signal is computed for each voxel by summing the number of signal-positive pixels in that voxel. Each image stack is registered in a multi-step process using both global affine and local deformable registration to the 3D Allen mouse brain reference atlas as previously described.^22,23^

###### Confocal imaging of rabies tracer-labeled starter cells

Antibody stained starter cells were scanned using a 10X objective lens and using a 4 μm step size on a Leica SP8 TCS confocal microscope using appropriately matched fluorescent filters. Images were auto-stitched from tiled scanning into tif image files and compiled into maximum intensity projection images for every section of the injection site. A cell counting algorithm was used to initially identify starter cells from the injection site. Following automated identification of starter cells, each section was then manually corrected using ImageJ.^24^

Each image containing the injection site was adjusted for brightness and false positive or false negative starter cells were corrected using the Cell Counter tool and starter cells were assigned to cortical layers based upon Dapi staining patterns.

#### Data Production at the Allen Institute, HUST, and SEU-Allen Joint Center: Single Neuron Reconstructions

##### Animal subjects at the Allen Institute

Male and female transgenic mice at an average age of P56 were utilized for all experiments (viral tracer and single neuron reconstructions). Mice had ad libitum access to food and water and were group-housed (3-5 per cage) within temperature (21-22°C), humidity (40% Rh), and light (12hr: 12hr light/dark cycle) controlled rooms within the Allen Institute vivarium. Experiments involving mice were approved by the Institutional Animal Care and Use Committees of the Allen Institute for Brain Science in accordance with NIH guidelines. Cre lines crosses to reporter lines are listed in **Extended Data Table 5**, and include drivers: Gnb4-IRES2-CreERT2, Fezf2-CreER, Cux2-CreERT2, Pvalb-T2A-CreERT2, Sst-Cre, and Cre-dependent EGFP reporters: Ai139 or Ai166.^25^ Induction of CreERT2 driver lines was done by administration via oral gavage of tamoxifen (50 mg/ml in corn oil) at original (0.2 mg/g body weight) or reduced dose for one day in an adult mouse. The dosage for mice age P7-P15 is 0.04 ml. Mice were transcardially perfused with fixative and brains collected after >2 weeks post-tamoxifen dosing.

##### Imaging and post-acquisition processing at HUST

All tissue preparation has been described previously.^26^ Following fixation, each intact brain was rinsed three times (6 h for two washes and 12 h for the third wash) at 4°C in a 0.01 M PBS solution (Sigma-Aldrich Inc., St. Louis, US). Then the brain was subsequently dehydrated via immersion in a graded series of ethanol mixtures (50%, 70%, and 95% (vol/vol) ethanol solutions in distilled water) and the absolute ethanol solution three times for 2 h each at 4°C. After dehydration, the whole brain was impregnated with Lowicryl HM20 Resin Kits (Electron Microscopy Sciences Cat#14340) by sequential immersions in 50, 75, 100 and 100% embedding medium in ethanol, 2 h each for the first three solutions and 72 h for the final solution. Finally, each whole brain was embedded in a gelatin capsule that had been filled with HM20 and polymerized at 50°C for 24 h.

Whole brain imaging is realized using a fluorescence microscopic optical sectioning tomography (fMOST) system. The basic structure of the imaging system is the combination of a wide-field upright epi-fluorescence microscopy with a mechanic sectioning system. This system runs in a wide-field block-face mode but updated with a new principle to get better image contrast and speed and thus enables high throughput imaging of the fluorescence protein labeled sample (manuscript in preparation). Each time we do a block-face fluorescence imaging across the whole coronal plane (X-Y axes), then remove the top layer (Z axis) by a diamond knife, and then expose next layer, and image again. The thickness of each layer is 1.0 micron. In each layer imaging, we used a strip scanning (X axis) model combined with a montage in Y axis to cover the whole coronal plane (Li et al., 2010). The fluorescence, collected using a microscope objective, passes a bandpass filter and is recorded with a TDI-CCD camera. We repeat these procedures across the whole sample volume to get the required dataset.

The objective used is 40X WI with numerical aperture (NA) 0.8 to provide a designed optical resolution (at 520 nm) of 0.35 μm in XY axes. The imaging gives a sample voxel of 0.35 x 0.35 x 1.0 μm to provide proper resolution to trace the neural process. The voxel size can be varied upon difference objective. Other imaging parameters for GFP imaging include an excitation wavelength of 488 nm, and emission filter with passing band 510-550 nm.

##### Full neuronal morphology reconstruction at Allen and SEU-Allen

Vaa3D, an open-source, cross-platform visualization and analysis system, is utilized for reconstructing neuronal morphologies as described in detail recently.^25^ Critical modules were developed and incorporated into Vaa3D for efficient handling of the whole-mouse brain fMOST imaging data, i.e., TeraFly and TeraVR (ref). TeraFly supports visualization and annotation of multidimensional imaging data with virtually unlimited scales. The user can flexibly choose to work at a specific region of interest (ROI) with desired level of detail. The out-of-core data management of TeraFly allows the software to smoothly deal with terabyte-scale of data even on a portable workstation with normal RAM size. Driven by virtual reality (VR) technologies, TeraVR is an annotation tool for immersive neuron reconstruction that has been proved to be critical for achieving precision and efficiency in morphology data production. It creates stereo visualization for image volumes and reconstructions and offers an intuitive interface for the user to interact with such data. TeraVR excels at handling various challenging yet constantly encountered data situations during whole-brain reconstruction, such as noisy, complicated, or weakly-labeled axons.

Trained reconstructors use the Vaa3D suite of tools to complete their reconstructions. Completion is determined typically when all ends have very well labeled, enlarged boutons. A final QC-checking procedure is always performed by at least one more experienced annotator using TeraVR who reviews the entire reconstruction of a neuron at high magnification paying special attention to the proximal axonal part or a main axonal trunk of an axon cluster where axonal collaterals often emerge and branches are more frequently missed due to the local image environment being composed of crowded high contrasting structures. To finalize the reconstruction, an auto-refinement step fits the tracing to the center of fluorescent signals. The final reconstruction file (.swc) is a single tree without breaks, loops, or multiple branches from a single point..

##### Registration of fMOST imaged brains to Allen CCFv3

We performed 3D registration of each fMOST image series (i.e., the subject) to the CCFv3 average template (i.e., the target)^23^ using the following steps^25^: 1) fMOST images were down-sampled by 64×64×16 (X, Y, Z) to roughly match the size of the target brain, 2) 2D stripe-removal was performed using frequency notch filters, 3) Approximately 12 matching landmark pairs between subject and target were manually added to ensure correct affine transformation that approximately aligned the orientation and scales, 4) Affine transformation was applied to minimize the sum of squared difference (SSD) of intensity between target and subject images, 5) Intensity was normalized by matching the local average intensity of subject image to that of target image, 6) A candidate list of landmarks across CCF space was generated by grid search (grid size=16 pixels), and finally 7) our software searched corresponding landmarks in the subject image and performed local alignment.

#### Data Analyses at the Allen Institute

##### Quantification of whole brain anterograde projections from MOp-ul

We generated a weighted connectivity matrix with data obtained from all anterograde tracer experiments^19^ for **Figure 4**. Experiments and data are provided in **Extended Data Table 3**. Segmentation and registration outputs are combined to quantify signal for every voxel in CCFv3. To quantify signal per brain structure, segmentation results are combined for all voxels with the same structure annotation. We defined connection weight in these analyses as the fraction of total axon volume; *i.e*., the axon volume segmented per each brain region divided by the total axon volume across all regions, excluding the injection site (MOp). We note that even with stringent QC, informatically-derived measures of connection weights can include artifacts (false positives), and the AAV-EGFP tracer reports signal from labeled axons, including passing fibers and synaptic terminals. For this reason, all targets (n=628 total, 314 per hemisphere) were visually inspected for presence of axon terminals, and a binary mask generated to reflect “true positives” for these regions. We applied the true positive binary mask to remove true negative connections and regions with only fibers of passage. We compiled a weighted matrix and performed comparative analyses across tracer datasets acquired from multiple labs (Allen, Huang, Dong). In the case of data from the Huang lab, integration was straightforward as these experiments were directly registered to CCFv3 as in the Allen pipeline. The Dong lab data were mapped to CCFv3 by matching structure name. As the ontology of the CCFv3 is derived from the ARA, corresponding structures were easily identified for most regions.

##### Quantification of whole brain retrograde inputs to MOp-ul

We generated a weighted connectivity matrix with data obtained from all retrograde tracer experiments (Yao et al, manuscript in preparation) for **Figure 5**. Experiments and data are provided in **Extended Data Table 4**. The total volume of detected signal was informatically-derived for each brain structure in CCFv3, as described above for axon segmentation. In contrast to the heavily manual QC for axonal projection false positives, we estimated segmentation false positives per CCFv3 structure for the rabies data by quantifying segmentation results from n=89-97 “blank” brains; *i.e*., brains processed through the imaging and informatics pipeline without rabies-mediated GFP expression. The distribution of false positives per structure was used to set a minimum threshold of 6 standard deviations from the mean. Any structure not passing this threshold was set to “0”. Following this threshold step, the input connection weights were defined as the fraction of fluorescent signal segmented per brain region divided by the total volume above threshold for this set of regions, again excluding the injection site (MOp).

##### Quantification of whole brain single neuron projections from MOp-ul

We generated a weighted connectivity matrix with data obtained from all single neuron full morphology reconstruction experiments for **Figure 7**. Experiments and data are provided in **Extended Data Table 5**. Reconstruction and registration outputs were again combined to quantify axon reconstructed for every CCF voxel, and combined for all voxels within the same CCF structure to generate total axon volume per brain structure for each single reconstructed cell. For **Figure 7**, we summed voxels from the same structure across hemispheres to match the data format obtained for MouseLight MOp reconstructions, then calculated the fraction of total signal per structure.

##### Clustering analyses based on connection weights

Unsupervised hierarchical clustering was conducted with the online software, Morpheus, (https://software.broadinstitute.org/morpheus/). Proximity between clusters was computed using complete linkages with spearman rank correlations as the distance metric. The clustering algorithm works agglomeratively: initially assigning each sample to its own cluster and iteratively merging the most proximal pair of clusters until finally all the clusters have been merged. The software program GraphPad Prism was used for statistical tests and generation of graphs.

#### Data availability statement from the Allen Institute

Viral tracing. Most anterograde tracing data (including high resolution STPT images, segmentation, registration to CCFv3, and automated quantification of injection size, location, and distribution across brain structures) are available through the Allen Mouse Brain Connectivity Atlas portal (http://connectivity.brain-map.org/). When available, direct links are provided in **Extended Data Table 3** on the metadata tab. For both AAV and transsynaptic rabies viral tracing, we also provide links to access 3D CCF-registered data files (http://download.alleninstitute.org/publications/), to download original images through the Brain Image Library (https://www.brainimagelibrary.org/). These links can be found on the metadata tabs in **Extended Data Tables 3** and **4**.

Single cell reconstructions. Original fMOST image datasets are available to download through the Brain Image Library (https://www.brainimagelibrary.org/). Links to access the final reconstruction files ((http://download.alleninstitute.org/publications/, with and without registration to CCF) are also provided in **Extended Data Table 5** on the metadata tab.

#### Experimental strategies for genetic targeting of cortical pyramidal neurons (PyNs) lines to produce gene expression, cell-type specific input and output whole-brain imaging datasets

Cell distribution and anatomical tracer data was generated as part of the Comprehensive Center for Mouse Brain Cell Atlas within the Huang lab at Cold Spring Harbor Laboratory (CSHL). Experimental methods and procedures have been described previously^27–29^.

##### Gene knockin mouse driver mice

Both male and female mice ≥ P56 were utilized for all experiments. Mice had ad libitum access to food and water and were group-housed within a temperature-(21-22°C), humidity- (40% Rh), and light- (12hr: 12hr light/dark cycle) controlled room within the CSHL Laboratory Animal Resources. All experimental procedures were approved by the Institutional Animal Care and Use Committee (IACUC) of CSHL in accordance with NIH guidelines. Mouse knockin driver lines are deposited at Jackson Laboratory for wide distribution.

Knockin mouse lines PlexinD1-2A-CreER, Fezf2-2A-CreER, Tle4-2A-CreER were generated (Matho et al BioRxiv). Foxp2-IRES-Cre was generated by R. Palmiter. We crossed CreER drivers (PlexinD1-2A-CreER, Fezf2-2A-CreER, Tle4-2A-CreER) with reporter mice expressing nuclear GFP or tdTomato (R26-CAG-LoxP-STOP-LoxP-H2B-GFP or R26-CAG-LoxP-STOP-LoxP-tdTomato, Ai14) for cell distribution data collection.

For both cell distribution and anterograde tracing analysis, these mice were induced with a 100 mg•kg-1 dose of Tamoxifen (T5648, Sigma) dissolved in corn oil (20 mg•ml-1), administered by intraperitoneal injection at the appropriate age to allow for temporal control of the CreER driver. In the case of the Foxp2-IRES-Cre line, cell distribution data was acquired based on a systemic AAV injection of AAV9-CAG-DIO-EGFP (UNC Viral Core) diluted in PBS (5×10^11^ vg/mouse), injected through the lateral tail vein at 4 weeks of age with 100 μl total volume. Cell distribution datasets from Matho et al BioRxiv were analyzed in MOp region. Experiments are detailed in Matho et al BioRxiv.

##### Tracer injection experiments

For anterograde tracing, adeno-associated viruses (AAVs) serotype 8 (UNC Vector Core, Salk) were delivered by stereotaxic injection. Briefly, cell-type specific anterograde tracing was conducted in the mouse knockin CreER and Cre driver lines. CreER drivers were crossed with the Rosa26-CAG-LSL-Flp mouse converter line such that tamoxifen induction of CreER expression at a given time is converted to constitutive Flp expression for anterograde tracing with a Flp-dependent AAV vector. For anterograde tracing from Foxp2-IRES-Cre driver line, we used a Cre-dependent AAV to express EGFP in labeled axons. Three weeks postinjection, mice were transcardially perfused with 0.9% saline, followed by icecold 4% PFA in PBS, brains were dissected out and processed for tissue collection.

##### Input tracing

For cell-type specific mono-trans-synaptic rabies tracing of inputs, in animals aged approximately 1 month, a Cre-dependent starter virus expressing TVA, EGFP and the G protein was delivered in MOp-ul, followed three weeks later, by the enVA-pseudotyped G-deleted rabies virus, all administered with a pulled glass pipette as specified below. In the case of CreER drivers, the starter virus injection was followed by Tamoxifen induction two and seven days post-injection. Seven to 10 days post-injection of the mono-trans-synaptic rabies virus, mice were transcardially perfused with 0.9% saline, followed by icecold 4% PFA in PBS, brains were dissected out and processed for tissue collection.

##### Stereotaxic surgeries

###### Anterograde and input tracing

Adult mice were anesthetized by inhalation of 2% isofluorane delivered with a constant air flow (0.4 L•min-1). Ketoprofen (5 mg•kg-1) and dexamethasone (0.5 mg•kg-1) were administered subcutaneously as preemptive analgesia and to prevent brain edema, respectively, prior to surgery, and lidocaine (2-4 mg•kg-1) was applied intra-incisionally. Mice were mounted in a stereotaxic headframe (Kopf Instruments, 940 series or Leica Biosystems, Angle Two). Stereotactic coordinates of MOp-ul were identified. An incision was made over the scalp, a small burr hole drilled in the skull and brain surface exposed.

A pulled glass pipette tip of 20–30 μm containing the viral suspension was lowered into the brain; a 300-400 nl volume was delivered at a rate of 30 nl•min-1 using a Picospritzer (General Valve Corp); the pipette remained in place for 10 min preventing backflow, prior to retraction, after which the incision was closed with 5/0 nylon suture thread (Ethilon Nylon Suture, Ethicon Inc. Germany) or Tissueglue (3M Vetbond), and animals were kept warm on a heating pad until complete recovery. Detailed description of virus used can be found in Table **“MOp experiments across labs”**.

##### Histology and immunohistochemical processing

###### Input tracing

For cell-type specific mono-trans-synaptic rabies tracing of inputs, postnatal mice aged 2 months were anesthetized using Avertin and intracardially perfused with saline followed by 4% PFA in PBS; brains were post-fixed in 4% PFA overnight at 4 °C and subsequently rinsed three times, embedded in 3% agarose-PBS and cut 50–100 μm in thickness using a vibrating microtome (Leica, VT100S). Sections were placed in blocking solution containing 10% Normal Goat Serum (NGS) and 0.1% Triton-X100 in PBS1X for 1 hr, then incubated overnight at 4 °C with primary antibodies diluted blocking solution. Anti-GFP (1:1000, Aves, GFP-1020); anti-RFP (1:1000, Rockland Pharmaceuticals, 600-401-379) were used. Sections were rinsed 3 times in PBS and incubated for 1 h at room temperature with corresponding secondary antibodies (1:500, Life Technologies). Sections were washed three times with PBS and incubated with DAPI for 5 min (1:5,000 in PBS, Life Technologies, 33342) to stain nuclei. Sections were dry-mounted on slides using Vectashield (Vector Labs, H1000) or Fluoromount (Sigma, F4680) mounting medium.

#### Cell type specific whole-brain imaging of soma distribution, input and output maps with focus on MOp-ul

##### Whole-brain Serial Two Photon Tomography

We used the whole-brain STP tomography (TissueCyte 1000, TissueVision Inc. Somerville, MA) pipeline (perfusion, postfixation, agarose embedding, crosslinking, imaging at high x-y resolution (1 μm x 1 μm) every 50 μm along the rostrocaudal z-axis, stitching, registration and segmentation of cell bodies and axon projections) as described by Osten lab^5,8^.

Images underwent QC, followed by signal detection and registration to the Allen Mouse Brain Common Coordinate Framework, version 3 (CCFv3). Registration brain-wide datasets to the Allen reference Common Coordinate Framework (CCF) version 3 was performed by 3D affine registration followed by a 3D B-spline registration using Elastix software^10^, according to parameters established by Ragan et al 2012^30^ and Kim et al., 2015^31^. For axon projection analysis, we registered the CCFv3 to each dataset so as to report pixels from axon segmentation in each brain structure without warping the imaging channel. Image processing on fields of view included in figures was completed using Adobe/Photoshop software with linear level and nonlinear curve adjustments applied only to entire images.

##### Cell body from whole-brain STP data

PyN somata were automatically detected from cell-type specific reporter lines (R26-LSL-GFP or Ai14) by a convolutional network trained as previously described^5^.

##### Axon detection from whole-brain STP data

For axon projection mapping, PyN axon signal based on cell-type specific viral expression of EGFP was filtered by applying a square root transformation, histogram matching to the original image, and median and Gaussian filtering using Fiji/ImageJ software so as to maximize signal detection while minimizing background auto-fluorescence. A normalized subtraction of the autofluorescent background channel was applied and the resulting thresholded images were converted to binary maps. 3D rendering was performed based on binarized axon projections and surfaces were determined based on the binary images using Imaris software (Bitplane). Projections were quantified as the fraction of pixels in each brain structure relative to each whole projection. We registered the CCFv3 to each dataset so as to report cells detected and pixels from axon segmentation in each brain structure without warping the imaging channel.

##### Microscopy imaging of cell-type specific input mapping

Imaging from serially mounted sections was performed using x5 objective on a Zeiss Axioimager M2 System equipped with MBF Neurolucida Software (MBF). To image starter cells, sections encompassing the injection site were imaged using a x20 objective with a 5 mm step size on a Zeiss LSM 780 or 710 confocal microscope (CSHL St. Giles Advanced Microscopy Center) using appropriately matched fluorescent filters. Images were auto-stitched from tiled scanning into tif image files and compiled into maximum intensity projection images for sections encompassing the injection site. Input cells were manually annotated within the serial sections to extract their position within the dataset. We matched the serial sections to the corresponding sections from CCFv3. Then, we placed fiduciary landmarks on both data and CCFv3 sections for warping conducted using Moving Least Squares in Fiji/ImageJ.

#### Cell type specific whole-brain image dataset presentation

Cell type specific anterograde viral tracing data generated (high resolution STPT images and registration to CCFv3) are available through the Mouse Brain Architecture Cell Type project (http://brainarchitecture.org/cell-type/projection). Cell type specific anterograde viral tracing, cell distribution and input tracing image datasets are available through the Brain Image Library (https://www.brainimagelibrary.org/). Cell distribution and anterograde tracing image datasets can also be viewed as imagesets registered to the Allen CCF by the Osten lab using Neuroglancer (https://github.com/google/neuroglancer). Links to these various portals can be found in the metadata tabs in **Extended Data Tables 3** and **4**.

### Giorgio Ascoli Lab

#### Data Analysis

##### Dendritic morphology analysis

Several consortium partners in this project contributed two neuronal reconstruction datasets (i.e., UCLA/USC and AIBS, Extended Data Figure 20). Both entailed sparse labeling of layer 2-5 pyramidal neurons using similar though distinct methodologies. The UCLA/USC contribution crossed Etv1-CreERT2 (layer 5-specific) and Cux2-CreERT2 (layers 2-4) mice with the Cre-dependent MORF3 (mononucleotide repeat frameshift) genetic sparse-labeling mouse line^32^. The MORF3 reporter mouse express a farnesylated V5 spaghetti monster fusion protein^33^ from the *Rosa26* locus when both the LoxP flanked transcriptional STOP sequence is removed by Cre and when stochastic-mononucleotide repeat frameshift occurs^34^. After perfusion, the tissue was cut into 500 μm-thick coronal slices, iDISCO+ cleared^9^ with a MORF-optimized protocol, stained with rabbit polyclonal anti-V5 antibody (1:500) followed by AlexaFluor 647-conjugated goat anti-rabbit secondary antibody (1:500) and NeuroTrace. Sections were imaged via a 30x silicone oil immersion lens with 1 μm z step on a DragonFly spinning disk confocal microscope (Andor). These tissue generation and processing methods are detailed in Veldman et al.^32^. Composite images of neurons were viewed with Imaris image software, manually reconstructed with Aivia reconstruction software (v.8.8.2, DRVision), and saved in the nonproprietary SWC digital morphology file format^35^.

The AIBS contribution crossed Cux2-CreERT2, Fezf2-CreER (layer 5-specific), and Pvalb-T2A-CreERT2 (layer 5) mice with the TIGRE-MORF (Ai166) fluorescent reporter line, which expresses farnesylated EGFP from the TIGRE locus^32^. Following tissue fixation, brains were processed using the fluorescence micro-optical tomography (fMOST) method. Briefly, brains were dehydrated in a graded ethanol series, immersed in a graded series of Lowicryl HM20 resin/ethanol solutions, and then polymerized in HM20 at 50°C. The crystalized brains were then block-face imaged in the coronal plane with the fMOST system, a combination microscope-microtome instrument that images the tissue surface, then shears off a 1 μm layer and images again, iteratively. These tissue generation and processing methods are detailed in Wang et al.^36^. Labeled neurons were reconstructed with Vaa3D software in a semi-automated, semi-user defined fashion^37^, utilizing the TeraFly and TeraVR modules enabling a virtual reality reconstruction environment, and reconstructions were saved as swc files.

Reconstructions from both datasets were analyzed concurrently at USC. Geometric processing of the reconstructions was performed with the Quantitative Imaging Toolkit (http://cabeen.io/qitwiki), allowing us to isolate the basal dendritic tree for analysis, and to render sample visualizations (Extended Data Figure 20A). The modified swc files were imported into NeuTube and morphometrics were obtained using L-Measure^38^. Since tissue preparation and data acquisition techniques can have significant effects on certain morphometric properties^39^, only measures that are insensitive to these effects were used in the present analyses. These measures were number of primary dendrites, remote bifurcation amplitude and tilt angles, branch order, branch path length, tortuosity, arbor depth, height, and width, Euclidian distance, total length, partition asymmetry, path distance, terminal degree, and terminal segments length. Data outputs were normalized by dividing all values within each dataset by the mean value of all layers 2-4 neurons for each morphometric. Principal component analysis was run on the data, and the first two components were graphed to create a low dimension scatterplot of the data (Extended Data Figure 20B). Wilcoxon signed rank tests were applied to all measures comprising the loadings for these two components, with the comparisons made between superficial (2-4) versus deep (5) layers (Extended Data Figure 20C); for the comparisons reported here the two datasets (AIBS and USC/UCLA) were not pooled together. A Sholl-like analysis was performed on the reconstructions to assess the distribution of dendritic distance as a function of relative path distance from the soma (Extended Data Figure 20D). Moreover, we carried out a comparative analysis of persistence diagram vectors^40^ of superficial vs. deep neurons for both datasets (Extended Data Figure 21E).

The Janelia MouseLight data set currently provides reconstructions from over 1000 neurons distributed across the entire rat brain. One can use the regions targeted by the neurons to distinguish and classify them. As case in point, the neurons of the primary motor cortex (MOp) may be analyzed in this manner. Before directly comparing the MOp neurons, some preprocessing of the data set is required. The MouseLight data set provides a point-by-point reporting of the regions targeted by each neuron. To distinguish between regions that are fully targeted from those that merely have axons passing through them, we thresholded the targeting counts in each region at 10, which yielded a final tally of 50 targeted regions distributed across the entire brain. Exclusive-or (XOR) pairwise comparisons are used to quantify the projection differences between two neurons. The targeted regions are then fully shuffled to produce a randomized distribution consistent with the regional projection patterns, corresponding to the ‘null’ hypothesis of continuous targeting patterns at the single cell level. The distribution of pairwise XOR distances of the shuffled data is then contrasted with the real pairwise distribution, which allows one to discern how much of the real distribution is accounted for by chance. To this end, given the non-normality of these distributions, we performed a 1-tail Levene test (Levene, 1960) to ascertain whether the variance of the experimental distribution was significantly larger than that of the shuffled distribution.

To estimate the relative proportions of the 6 clusters we matched their respective singlecell axonal patterns against the regional patterns from PHA-L anterograde tracing across all target regions. Specifically, the problem is equivalent to a set of constrained, weighted, linear equations that can be solved numerically by standard non-negative least-square (NNLS) or bounded-variable least-squares (BVLS) optimization. The NNLS algorithm solves the linear least squares problem arg minx ||Ax - b||2 with the constraint x ≥ 0^41^. The BVLS variant^42^ minimizes the same objective function, but subject to explicit boundary conditions. We used the respective R implementations nnls^43^ and bvls^44^. Boundary conditions for bvls were 0.01 for lower bound and 1 for upper bound. The results were consistent between the two methods.

#### Fluorescence micro-optical sectioning tomography (fMOST) methodology

##### Animal subjects

PlexinD1-2A-CreER, Fezf2-2A-CreER were generated by Josh Huang’s lab^29^, and were crossed with Rosa26-loxp-stop-loxp-flpo mice. All mice are kept in a suitable environment with sufficient water and food, and animal experiments are conducted in accordance with the requirements of the Animal Ethics Committee of Huazhong University of Science and Technology.

We used adult double-positive hybrid mice aged 2-3 months for experiments. After deep anesthesia (mixed with 2% chloral hydrate and 10% urantan), we cut scalp to expose the skull, 50nl of flp-dependent pAAV-EF1a-fDIO-TVA-GFP virus (8×1012 gc/ml, the UNC Vector Core) was injected in the motor cortex. Three days later, the mice were induced intraperitoneally with a low amount of tamoxifen (T5648, Sigma, dissolved in corn oil, diluted at a concentration of 5mg / ml, and the injection dose per mouse was 10g / 1ml), and then the virus expressed in brain for 5 weeks.

##### Perfusion and Resin Embedding

Mice were anesthetized with anesthetic, and perfused with PBS and 4% paraformaldehyde (PFA, Sigma-Aldrich Inc., St Louis, MO, USA). The mouse brain was peeled off and placed overnight in PFA, and then rinsed with PBS. Those samples were dehydrated in gradient alcohol and embedded into HM20. All dehydration and infiltration procedures are processed at 4 degrees.

#### Data Collection

##### Imaging and post-acquisition processing

The whole brain imaging is realized using a fluorescence micro-optical sectioning tomography (fMOST) system. The basic structure of the imaging system is the combination of a line scanning upright epi-fluorescence microscopy with a mechanic sectioning system. This system runs in a line scanning block-face imaging mode but updated with a new principle to get better optical sectioning image contrast and data acquisition speed and thus enables high throughput imaging of the fluorescence protein labeled sample (manuscript in preparation). Each time we do a block-face fluorescence imaging across the whole coronal plane (X-Y axes) in a thickness of 2.0 micron with an interval of 1.0 micron, then remove the imaged top layer (Z axis) by a diamond knife, and then expose next layer, and image again. In each layer imaging, we used a strip scanning (X axis) model combined with a montage in Y axis to cover the whole coronal plane. The fluorescence, collected using a microscope objective, passes a bandpass filter and is recorded with a sCMOS camera. We repeat these procedures across the whole sample volume to get the required dataset.

The objective used is 20X WI with numerical aperture (NA) 1.0 to provide a designed optical resolution (at 520 nm) of 0.35 μm in XY axes. The imaging gives a sample voxel of 0.35 x 0.35 x 1.0 μm to provide proper resolution to trace the neural process. The voxel size can be varied upon difference objective. Other imaging parameters for GFP imaging include an excitation wavelength of 488 nm, and emission filter with passing band 510-550 nm. Another excitation wavelength of 561 nm and emission filter with long passing band of 590 nm is used to acquire the cytoarchitecture reference by imaging the propidium iodide-stained (PI) cellular nuclei (Gong et al., 2016)

##### fMOST Data Analysis Pipeline (for Fig.7, sample No. 193377, 193663)

The fMOST datasets have two color channels. The green channel contains fluorescent protein signal from labeled neurons is used to reconstruct neuronal morphology. The red channel contains propidium-iodide(PI) signal with clear contours of most brain regions, is used to map original images to CCF. We have built a data analysis pipeline to perform neuron reconstruction and spatial mapping.

We used GTree software to reconstruct neuronal morphology with human-computer interaction^45^. GTree is an open-source GUI tool, it offers a special error-screening system for the fast localization of submicron errors and integrates some automated algorithms to significantly reduce manual interference. To random access image block from brain-wide datasets, the original image (green channel) was pre-formatted to TDat, an efficient 3D image format for terabyte- and petabyte-scale large volume image^40^. GTree has a plugin to import TDat formatted data, and save reconstructions with original position in SWC format. All reconstructions were performed back-to-back by experienced technician and checked by neuroanatomists.

We used BrainsMapi to complete the 3D registration^46^. Specifically, the image of red channel is down-sampled to an isotropic 10 μm resolution consistent with the CCF atlas. We conduct the registration by several key steps including the initial position correction, regional feature extraction, linear and nonlinear transformation and image warping. Among them, a set of anatomically invariant regional features are extracted manually by Amira (version 6.1.1; FEI, Mérignac Cedex, France) and automatically by DeepBrainSeg^47^. Based on these, the unwarping neuron reconstructions can be accurately transformed to CCF.

### BARseq

#### Data Production

##### Animal subjects

Eight-week old male mice were utilized for all experiments. Mice had ad libitum access to food and water and were group-housed within a temperature-(21-22°C), humidity- (40% Rh), and light- (12hr: 12hr light/dark cycle) controlled room within the CSHL Laboratory Animal Resources. All animal procedures were carried out in accordance with Institutional Animal Care and Use Committee protocol 19-16-13-10-07-03-00-4 at Cold Spring Harbor Laboratory.

##### Sindbis injection experiments

Two eight-week old C57BL/6J males (Jackson Laboratory) were injected in the motor cortex at 0.5 mm AP, 1.5 mm ML at 500 μm and 900 μm depths. At each depth, 150 nL of a JK100L2 Sindbis barcoded library (Chen et al., 2019; Kebschull et al., 2016) diluted 1:2 in PBS was injected.

#### Data Collection

##### BARseq and post-acquisition processing

After 24 hrs, each animal was sacrificed, and a 3 mm diameter biopsy punch was used to remove the area around the injection site. The injection site punch was mounted in OCT and snap-frozen in liquid nitrogen – isopentane bath. The rest of the brain was frozen on dry ice, cryo-sectioned to 300 μm slices, and processed for projection site sequencing.

The projection sites were dissected manually based on a list of targets designed to match bulk GFP tracing results. Several large brain areas, including the striatum, the thalamus, and the medulla, were dissected into multiple samples to obtain a relative position of the projections. A list of designed dissection sites and their coordinates is provided in Supp. Table S1. The images of the slices after dissection are provided at Mendeley data (see data availability). After dissection, RNA barcodes were extracted from the samples, amplified, and sequenced as described previously^48^. The raw sequencing data and processed barcode counts are available from SRA and Mendeley data (see data availability).

The injection site was cryo-sectioned to 20 μm sections and the sections were mounted onto Superfrost Plus Gold slides (Electron Microscopy Sciences) using NOA81 glue and a home-made tape-transfer system^49^. Six slides encompassing the center of the injection site was then processed for BARseq library preparation and sequencing as previously described^48^.

Sequencing images were captured on an Olympus IX81 microscope with a Crest X-light v2 spinning disk confocal, an 89north LDI 7-channel laser, and a Photometrics Prime BSI camera. Image acquisition was controlled through micro-manager^50^. Filters and lasers used for each imaging channel are provided in Supp. Table S2. At each sequencing cycle, a z-stack with a step size of 7 μm and a span of 50 μm centered on the injection site was taken using a UPLSAPO 10× NA 0.45 objective for all four sequencing channels and the DIC channel. After the last sequencing cycle is done, we performed DAPI staining and imaged a 3 × 3 tile with 15% overlap to roughly cover the whole punch.

To process the images, we first max-projected all z-stacks, followed by median filtering, channel bleed-through correction, and top-hat filtering. The pre-processed images, including the stitched image of the 3 ×3 tile, were then registered to the first sequencing cycle using ECC image alignment^51^. Cells were identified by first finding local maxima above a prominence threshold, then segmenting somata using the sequencing signals around each local maxima. The barcodes were base-called by calling the channel with the maximum signal in each sequencing cycle across all pixels within the segmented cells. We then manually marked the outline of the top and the bottom of the cortex, and the medial and lateral boundaries of the punch in the stitched 3 × 3 images. The distance of each cell to these four boundaries were determined to find the relative position of the cell within the cortex. The *in situ* barcodes were matched to the barcodes at the projection sites, allowing one mismatch but no ambiguous matches (i.e. an *in situ* barcode cannot match to two projection barcodes with equal hamming distance). Only *in situ* barcodes with a minimum quality score of at least 0.8 across all cycles were kept for further analysis.

Some barcodes were seen in more than one cells. This could be caused by a cell being sliced into two in adjacent slices, or the apical dendrite of a cell being segmented as a cell body by mistake, or by two cells being labeled with the same barcode. The latter two possibilities would either cause the cell to be identified in the wrong laminar position or a loss of single-cell resolution in projection mapping. We thus removed all pairs of cells with the same barcodes if they had a laminar difference of more than 100 μm or were not found on the same or adjacent slices. For the remaining pairs, we kept the cell in the deeper layer (because apical dendrites were likely more superficial than the somata). We further filtered the cells so that all cells have a strongest projection of at least 5 molecule counts. This resulted in 10,299 projection neurons for further analyses.

#### Data Analysis

Raw projection barcodes were first normalized by spike-in counts, and further normalized between the two brains so that neurons with non-zero counts in each projection area have the same mean across the two brains. We then performed hierarchical k-mean clustering on log transformed and spike-in corrected projection strengths to identify the major classes. However, this clustering failed to identify small clusters with distinct laminar positions. To find subclusters with distinct laminar distributions, we used a second clustering method based on binary projection patterns. From a population of neurons, we first split off one subcluster with a particular binary projection to up to three brain areas. For example, a subcluster can be defined as having projections to the contralateral primary motor cortex, the ipsilateral caudal striatum, but not the caudal medial section of the ipsilateral thalamus. These projections were chosen to maximize the reduction in the entropy of the laminar distribution of neurons. This process was then iterated over the two resulting subclusters, until no subclusters resulted in statistically significant reduction in entropy (p < 0.05 without multiple testing correction). This process resulted in many clusters, some of which may have similar laminar distributions. We then built a dendrogram based on the distance in projection space among the resulting clusters and iteratively combined subclusters similar in laminar distribution. Two subclusters were considered similar in laminae if differences in their laminar distributions were not statistically significant (p < 0.05 using rank sum test with Bonferroni correction) and their median laminar positions were within 200 μm. This process was iterated over each split, starting from ones between the closest leaves/branches. We stopped combining clusters at the level of major classes.

To compare BARseq dataset to single-cell tracing, we randomly down-sampled BARseq dataset to the same sample size as the single-cell tracing dataset. We further combined ipsilateral and contralateral cortical areas and combined all samples of the same non-isocortex brain divisions together. This resulted in an axonal resolution that can be compared to the single-cell tracing dataset. We then combined this down-sampled and low resolution BARseq dataset with the traced neurons and analyzed the joint dataset. T-SNE was performed in MATLAB. Clustering was performed using two layers of Louvain community detection^52^ in MATLAB.

Matching BARseq clusters to single-cell tracing clusters was done using the common axonal resolution, but full-size BARseq dataset using MetaNeighbor^53^. To test the homogeneity of clusters, we downsampled the datasets with replacement to different sizes (1000 random samples per cluster size) and calculated the correlation between the downsampled cluster centroids to the full-data cluster centroids.

#### Data Presentation

Raw bulk sequencing data are deposited at SRA (SRR12247894). Raw *in situ* sequencing images are deposited at Brain Image Library. Processed projection data and *in situ* sequencing data are available from Mendeley Data (preview available at https://data.mendeley.com/datasets/tmxd37fnmg/draft?a=320d5121-a325-413d-83fa-50c97b0db266).

#### Informatics Tools and Code availability

Processing scripts for *in situ* sequencing images, processed data, annotated BARseq dissection images, and analysis codes are available from Mendeley Data (preview available at https://data.mendeley.com/datasets/tmxd37fnmg/draft?a=320d5121-a325-413d-83fa-50c97b0db266).

### Nissl data set generation and automated MOp-SSp boundary detection

#### Data Production

##### Animal subjects

Ten wildtype male C57BL/6 mice (11-20 weeks, weight 20-26.4g) were involved in the cytoarchitectural analysis of the MOp-SSp border variation.

##### Histology and immunohistochemical processing

The entire brain was sectioned coronally with 20 μm thickness using the tape-transfer method to minimize tissue distortion^49^. In one brain, consecutive coronal sections were processed with Nissl staining for cytoarchitecture. In all other brains, alternative coronal sections were processed with Nissl staining.

#### Data Collection

##### Imaging and post-acquisition processing

Bright field imaging was performed using Nanozoomer 2.0 HT (Hamamatsu, Japan) with a 20x objective. Images of individual coronal sections were converted into 8-bit RGB images at 0.46 μm/pixel in-plane resolution in JPEG2000 format (Figure S1A).

#### Data Analysis

A diffeomorphic registration pipeline^54^ was applied to register the whole stack of coronal sections into 3D volume and mapped with the Allen Mouse Brain Atlas^55^. Two versions of atlas annotation were applied: Common Coordinate Framework (CCF) published in 2011^56^ and 2017^57^. For sections within a range of AP-1.5mm~+1.5mm, a thresholding method was used to eliminate the high-intensity pixels in the Red and Green channels and retain the full spectrum Blue channel intensities. The segmented Nissl stained cells in the whole section constitute a binary mask *M_N_* (Figure S1B). The binary cortical mask, *R_ctx_*, was extracted from the mapped CCF3 segmentation. A morphological thinning operation was applied on the boundary-smoothed *R_ctx_* to generate a smooth pseudo medial axis (*medAX_s_*) of the entire cortical band. A centroid *C_ctx_* was determined based on *R_ctx_*. The angle 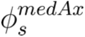 to each point sampled uniformly on *medAX_s_* was calculated clockwise from the *C_ctx_* to generate the parameterized representation of *medAX_s_* (Figure S1C).

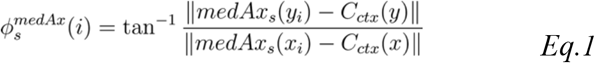

The cortical normal *N_medAx_* at each point on the *medAX_s_* was sampled with a uniformly distributed 1,000 points spanning the cortical depth, to determine the profile representation of the flattened cortex map. That is, for *N_medAx_* := {*x_i_*, *y_i_*},

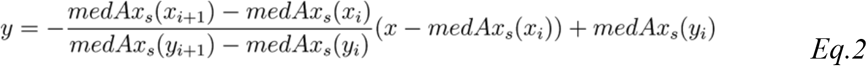

Figure S1D shows an example of cortical normals *N_med_A_x_* sampled at 1,000 cortical depths of the Nissl mask *M_N_* for all angles 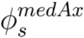. The density value of each point represents the density of segmented Nissl mask within a surrounding neighborhood of ~93×93μm^2^.

To identify the MOp border with cell-sparse SSp layer 5a band, underneath the thick and densely-packed granular cell layer 4, we took fractional change of the flattened cortical profile across cortical depth

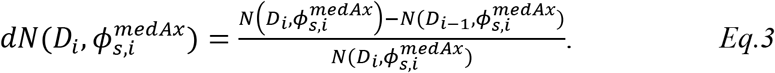

All local minima were identified within the middle 30%-70% of cortical depth in dN. The connected objects with dN<-ε, where 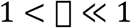, containing the most minima on either side of 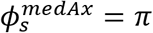, where retained as a putative layer 5a of SSp (Figure S3E). By aligning all profiles in the stack ranging from −1.5mm to +1.5mm from Bregma in AP direction, two smooth objects of putative SSp layer 5a were constructed in 3D (Figure S1F). The border *B_n_* with MOp was identified as *N_medAx_* at 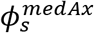 associated with the “inner bounds” of either object (close to 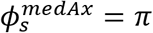) on each profile (red lines in Figure S1F). *B_n_* of all flattened cortical profiles were projected back to the registered brain space (curved cortex) and constructed a smooth surface ***S_B_*** (Figure S1G). By reverse transformation of atlas mapping, ***S**_B_* was projected into the common atlas space. The intersecting line between ***S**_B_* and the dorsal surface of the cortex was compared with that of CCF2 and CCF3 in the subsequent analysis (Figure 2f, ED Fig 5a).

#### Data Presentation

Cytoarchitecture data generated is published online as part of the Mouse Brain Architecture web portal (http://brainarchitecture.org/). Links to individual datasets are available at https://docs.google.com/spreadsheets/d/1MPJuzSD-vGTJyuaj5SBUx345_K7MShwWy0zInPAbwSc/edit?usp=sharing.

#### Informatics Tools and Code availability

All related code for border detection are published on Github (https://github.com/bingxinghuo/AutoSegmentBrain/).

##### Processing and imaging of MORF tissue

Several consortium partners in this project contributed two neuronal reconstruction datasets (i.e., UCLA/USC and AIBS, Figure 6). Both entailed sparse labeling of layer 2-5 pyramidal neurons using similar though distinct methodologies. The UCLA/USC contribution crossed Etv1-CreERT2 (layer 5-specific) and Cux2-CreERT2 (layers 2-4) mice with the Cre-dependent MORF3 (mononucleotide repeat frameshift) genetic sparse-labeling mouse line^32^. The MORF3 reporter mouse express a farnesylated V5 spaghetti monster fusion protein^33^ from the *Rosa26* locus when both the LoxP flanked transcriptional stop sequence is removed by Cre and when stochastic-mononucleotide repeat frameshift occurs^32^. After perfusion, the tissue was cut into 500 μm-thick coronal slices, iDISCO+ cleared^9^ with a MORF-optimized protocol, stained with rabbit polyclonal anti-V5 antibody (1:500) followed by AlexaFluor 647-conjugated goat anti-rabbit secondary antibody (1:500) and NeuroTrace. Sections were imaged via a 30x silicone oil immersion lens with 1 μm z step on a DragonFly spinning disk confocal microscope (Andor). These tissue generation and processing methods are detailed in Veldman et al.^32^. Composite images of neurons were viewed with Imaris image software, manually reconstructed with Aivia reconstruction software (v.8.8.2, DRVision), and saved in the non-proprietary SWC digital morphology file format^35^.

The AIBS contribution crossed Cux2-CreERT2, Fezf2-CreER (layer 5-specific), and Pvalb-T2A-CreERT2 (layer 5) mice with the TIGRE-MORF (Ai166) fluorescent reporter line, which expresses farnesylated EGFP from the TIGRE locus^32^. Following tissue fixation, brains were processed using the fluorescence micro-optical tomography (fMOST) method. Briefly, brains were dehydrated in a graded ethanol series, immersed in a graded series of Lowicryl HM20 resin/ethanol solutions, and then polymerized in HM20 at 50°C. The crystalized brains were then block-face imaged in the coronal plane with the fMOST system, a combination microscope-microtome instrument that images the tissue surface, then shears off a 1 μm layer and images again, iteratively. These tissue generation and processing methods are detailed in Wang et al.^36^. Labeled neurons were reconstructed with Vaa3D software in a semi-automated, semi-user defined fashion^37^, utilizing the TeraFly and TeraVR modules enabling a virtual reality reconstruction environment, and reconstructions were saved as swc files.

##### Dendritic morphology analysis

Reconstructions from both datasets were analyzed concurrently at USC. Geometric processing of the reconstructions was performed with the Quantitative Imaging Toolkit (http://cabeen.io/qitwiki), allowing us to isolate the basal dendritic tree for analysis, and to render sample visualizations (Figure 6A). The modified swc files were imported into NeuTube and morphometrics were obtained using L-Measure^38^. Since tissue preparation and data acquisition techniques can have significant effects on certain morphometric properties^39^, only measures that are insensitive to these effects were used in the present analyses. These measures were number of primary dendrites, remote bifurcation amplitude and tilt angles, branch order, branch path length, tortuosity, arbor depth, height, and width, Euclidian distance, total length, partition asymmetry, path distance, terminal degree, and terminal segments length. Data outputs were normalized by dividing all values within each dataset by the mean value of all layers 2-4 neurons for each morphometric. Principal component analysis was run on the data, and the first two components were graphed to create a low dimension scatterplot of the data (Figure 6B). Wilcoxon signed rank tests were applied to all measures comprising the loadings for these two components, with the comparisons made between superficial (2-4) versus deep (5) layers (Figure 6C); for the comparisons reported here the two datasets (AIBS and USC/UCLA) were not pooled together. A Sholl-like analysis was performed on the reconstructions to assess the distribution of dendritic distance as a function of relative path distance from the soma (Extended data Fig. 21). Moreover, we carried out a comparative analysis of persistence diagram vectors^40^ of superficial vs. deep neurons for both datasets (Extended Data Figure 21E).

### eFLASH-based 3D immunohistochemistry from Chung lab

#### Data Production

##### Animal subjects

Young adult male mice (C57BL/6) were purchased from Jackson (Stock No. 000664) and were housed in a 12 hr light/dark cycle with unrestricted access to food and water. All experimental protocols were approved by the MIT Institutional Animal Care and Use Committee and the Division of Comparative Medicine and were in accordance with the guidelines from the National Institute of Health.

##### Histology and immunohistochemical processing

Mouse brains were preserved using SHIELD technology.^58^ Briefly, mice were transcardially perfused at PND 57 with ice-cold 1X PBS and then with SHIELD perfusion solution (10% (w/v) P3PE and 4% PFA (w/v) in 1X PBS). The brains were dissected and incubated in the SHIELD perfusion solution at 4 °C for 48 h, and then transferred to the SHIELD-OFF solution (1X PBS containing 10% (w/v) P3PE). After incubation at 4 °C for 24 h, the brains were placed in the SHIELD-ON solution (0.1 M sodium carbonate buffer at pH 10) prewarmed to 37 °C and then incubated at 37 °C for 24 h. The brains were then washed in 1X PBS with 0.02% sodium azide at room temperature for O/N and were rapidly cleared using stochastic electrotransport (SmartClear Pro, LifeCanvas Technologies).

SHIELD-processed brains were immunostained using eFLASH^59^. A Neurofilament-M antibody (MCA-3H11, Encor Biotechnology) and NeuN antibody (266004, Synaptic Systems) were used to immunostain a whole mouse brain. After staining, brains were washed in 1X PBS with 0.02% (w/v) sodium azide at RT overnight, and were fixed with 4% (w/v) PFA solution in 1X PBS at RT overnight to prevent the dissociation of bound antibodies. After washing with 1X PBS with 0.02% (w/v) sodium azide at RT with multiple solution exchanges, brains were optically cleared.

Protos-based immersion medium was used for optical clearing of whole brains^58^. Brains were firstly incubated in half-step solution (50/50 mix of 2X PBS and the Protos-based immersion medium) at 37°C overnight. Afterwards, the samples were moved to the pure immersion medium and incubated at 37°C overnight.

#### Data Collection

##### Imaging and post-acquisition processing

Optically cleared brains were imaged with an axially-swept light-sheet microscope (SmartSPIM, Lifecanvas Technologies). Imaging was done using 3.6x objective (custom Lifecanvas design; 0.2NA, 12mm working distance, 1.8um lateral resolution) and using three lasers (488um, 561um, and 642um wavelengths). Acquired data was post-processed and region-segmented as described in Swaney et al.^60^ Acquired 3D images were aligned to the Allen brain reference atlas (CCF V3) based on tissue autofluorescence from 488nm laser illumination. Delineation of cortical layers in MOp was refined using anti-NeuN and anti-Neurofilament-M signals. Calculations and visualizations were done using Imaris (Bitplane) and Nuggt python package^60^.

